# Multi-Tissue Profiling Reveals tissue-specific protein regulation and relationships Between Protein Quantitative Trait Loci (pQTLs) and Cardiometabolic Disease

**DOI:** 10.1101/2025.06.30.662294

**Authors:** April E Hartley, Katyayani Sukhavasi, Sile Hu, Matthew Traylor, Mar Gonzalez-Ramirez, Kristian Ebbesen Hanghøj, Husain Talukdar, Arno Ruusalepp, Ellen Björkegren, Johan LM Björkegren, Joanna MM Howson, Yalda Jamshidi

## Abstract

Integrating genetic data with protein levels, known as protein quantitative trait loci (pQTLs), can enhance our understanding of disease mechanisms and provide actionable insights for drug discovery, by guiding the direction of therapeutic interventions, clarifying mechanisms of action, and predicting potential side effects. However, most pQTL studies have focused on the plasma proteome, overlooking tissue-specific effects.

Here, we investigate the plasma and tissue proteome and derive tissue-specific pQTLs in a unique dataset derived from a cohort of 284 STARNET patients, predominantly male, with a mean age of 65 years and a high prevalence of coronary artery disease (CAD). Importantly, our dataset includes paired tissue samples from aortic wall, mammary artery, liver, and skeletal muscle alongside plasma, allowing for a comprehensive comparative analysis across tissues—all from the same individuals. We employed the Olink Explore 3.2k platform to assess relative protein levels in each tissue. We identify 608 *cis-*pQTLs, the majority of which are found in plasma, reflecting greater protein variability. Notably, we find 13 proteins with exclusive tissue-specific pQTLs, underscoring distinct as well as shared genetic influences across tissues. Colocalization analyses reveal shared genetic regulation between tissue proteins and cardiometabolic traits, including LDL, HDL, and triglycerides levels, implicating proteins such as PNLIPRP2, SORT1, and PRSS53 as potential mediators of lipid regulation. Furthermore, Mendelian randomization analyses suggest a liver-specific role for SORT1 and PSRC1 in modulating CAD risk and lipid profiles.

Our findings highlight the importance of profiling tissue-shared, and tissue-specific, protein expression and pQTLs to elucidate disease mechanisms and accelerate precision drug and biomarker discovery.

## Introduction

Proteins serve as crucial intermediaries between genes and diseases, directly participating in biological processes, and making their measurement essential for the identification of novel drug targets (1). Drug targets supported by genetic evidence are more than twice as likely to succeed in clinical trials (2–5). Nevertheless, there are thousands of associations across the genome and the function of many are not well understood, nor the molecular effects of a therapeutic perturbation, including tissue of action. Integrating genetic data with protein levels, specifically to identify genetic variants strongly associated with protein levels (known as protein quantitative trait loci or pQTLs), enhances causal inference for the role of specific proteins in disease (6). This approach can reveal mechanisms through which a drug may act to prevent or treat disease.

In recent years, there has been a significant increase in the identification of pQTLs for protein levels in plasma. Notably, pQTLs were identified for 2,415 proteins measured in ∼50,000 UK Biobank participants (7). This progress has been aided by the development of affinity-based assays capable of quantifying thousands of proteins in a single biological sample (8). Despite these advancements, the vast majority of pQTL studies have focused on the plasma proteome, missing insights on the source of tissue-specific proteins that do not circulate in blood (1,9,10). This limitation poses a challenge in assessing the role of these tissue-specific proteins in disease, as their pQTLs may not be identified in plasma samples. Additionally, using plasma protein levels to estimate drug target effects could also lead to incorrect decisions on drug target mechanism of action if pQTL effects differ in direction or magnitude between tissue of origin and plasma.

To address this gap, we performed antibody-based proteomic profiling using Olink technology (8) from donors across four key cardiometabolic tissues (Liver, Skeletal Muscle, Aorta and Mammary Artery), in a total of 1,242 samples, isolated from 284 patients in the Stockholm-Tartu Atherosclerosis Reverse Engineering Task (STARNET) cohort (11,12). Our study is the first to generate pQTLs in multiple cardiometabolic tissues sampled from the same individuals, providing a unique dataset that links genetic variation to protein levels in disease-relevant tissues. This tissue-based approach allows us to identify proteins with mechanisms specific to certain tissues, which are crucial for understanding disease pathways and developing targeted therapies.

## Methods

### The STARNET Cohort

Coronary artery disease (CAD) patients eligible for coronary artery by-pass grafting (CABG) surgery (n=200) and controls eligible for other open-heart surgeries (e.g. valve replacement, n=84), verified to be CAD-free in pre-operative coronary angiograms, were enrolled at the Department of Cardiac Surgery, Tartu University Hospital. Informed consent was obtained from all subjects (Ethics Review Committee on Human Research of the University of Tartu, IRB no:289/T-12). Each enrolled participant completed a questionnaire to assess disease history, current drug regimens, and lifestyle. Inclusion criteria were patients eligible for heart surgery. Exclusion criteria were other severe systemic disease such as active cancer or inflammatory disease. During open-heart surgery, atherosclerotic aortic arterial wall (AOR), mammary artery (MAM, only in individuals with CAD), liver (LIV) and skeletal muscle (SKM) biopsies were obtained and immediately preserved in Allprotect Tissue Reagent (Qiagen) and frozen at −80°C. Preoperative blood samples were collected for biochemical screens, plasma, whole blood RNA and DNA isolations.

### DNA extraction, genotyping and quality control

DNA was isolated from whole blood stored in PAXgene blood DNA tube (QIAGEN) using the PAXgene blood DNA kit (PreAnalytiX). DNA qualities were assessed with the NanoDrop 2000 spectrophotometer (Thermo fisher scientific). The blood DNA samples were genotyped at the Core Facility of Genomics, Institute of Genomics, University of Tartu, using the Illumina Infinium Global Screening Array-24 v3.0 (customized version GSAv3.0_EST) following the standard protocol. Illumina GenomeStudio 2.0.5 was used for initial quality control (QC) to check call rates and overall performance. As additional sample QC, pairs of sample duplicates and samples with more than 3% missing calls were excluded. Genetic variants with more than 3% missingness or deviating from HWE (p-value <10^−7^) were excluded from analyses. Following variant normalization, allele frequencies were matched against gnomAD (https://gnomad.broadinstitute.org/) allele frequencies to identify ambiguous alleles and large allele frequency deviations. After QC, genotype data was imputed to TOPMed version 3 (13) using the TOPMed imputation server following the recommended data preparation steps (14).

### Proteomic profiling of plasma and tissue samples

Plasma was isolated from EDTA blood tubes by centrifugation for 10 minutes (RT) 300 rcf (acceleration 4, deceleration 0). The plasma (supernatant) was collected in 2ml Eppendorf tubes and frozen at -80°C. Tissue protein isolation was performed using a 10x RIPA buffer (Sigma, R0278) with protease inhibitor cocktail (cOmplete^™^, Mini Protease Inhibitor Cocktail, Roche). Tissue was weighed and buffer added with inhibitor in the ratio of 1:4 (i.e to 0.1g of tissue add 0.4ml of buffer with inhibitor). All steps were carried out at 2-8°C. Tissue was homogenized on ice using an Ultra-Turrax homogeniser until the tissue was properly minced, then incubated on ice on a shaker for 1 hour. Next, tissue was centrifuged at 5000rpm for 15mins at 4°C. The pellet was discarded and tissue protein isolated (supernatant) was measured using Bradford reagent (ThermoFisher) in a microplate reader.

Plasma and tissue proteomic analysis was performed using the Olink proximity extension assay and the Explore 3,072 panels. The methodology has been described in detail elsewhere (15). Briefly, two proximity probes were developed for each protein. These probes are an antibody linked to a unique complimentary oligonucleotide. When the two probes bind to the protein, they come into proximity and the oligonucleotides hybridize, which enables the DNA to be amplified and then quantified by next generation sequencing. The Explore 3,072 panel consists of eight panels of 384 proteins, and each panel has four dilution blocks to allow for different ranges of the target proteins. Three controls are added to each sample: an incubation control, which is a non-human assay used for quality control; the extension control, which is an antibody coupled to a unique pair of oligonucleotides in proximity and is used for normalization; and the amplification control, which is synthetic double stranded DNA and is used for quality control. Additionally, control samples are added to each plate for quality control and normalization: the biological sample control, which is run in duplicate on each plate to calculate intra- and inter-plate CVs (*Supplementary Table ST1*); a negative control sample is run in triplicate on each plate, which is buffer and is used to calculate limit of detection (5% of plasma proteins had a median value below the limit of detection, 4% for liver and aorta, 7% for muscle and 5% for mammary artery); and three plate controls are also added to each plate, which are pooled healthy plasma samples used for normalization.

Counts of each sequence are translated to Normalized Protein eXpression (NPX) values using Olink’s MyData Cloud Software. A plate-control normalized NPX value for each assay for each sample is calculated by subtracting the assay-specific median of the plate controls from the log_2_ ratio of assay counts per sample to the counts of the extension control. Further normalization across plates can be performed using intensity normalization, where the data are adjusted to have the same median value for an assay on all plates. For each assay, a median value is calculated for all samples and plates; a plate-specific median is then calculated for each assay, and this is subtracted from every value on the plate; for each assay, the overall median value is added. This normalization process assumes there is no difference expected between the median signal for an assay on one plate compared to another. We used the intensity normalized data for plasma as cases and controls were randomized across plates and we expected median assay values to be the same across plates. For tissue, we randomized tissues and cases *vs* controls across all plates, but due to differing proportions of tissues on each plate, we did not expect the median to be the same for an assay across all plates and therefore used the plate-controlled NPX values.

We removed 40 plasma proteins and 84 tissue proteins which did not meet the batch release quality control criteria defined by Olink. Outlier samples were identified and remover based on a PC1 or PC2 value more than five standard deviations from the mean, and a median NPX value across all assays more than five standard deviations from the mean of the median values for at least four of the eight Olink panels. This identified one plasma sample to remove. Four plasma samples were duplicated, therefore we took the mean value for each assay of each pair of duplicates.

Associations of protein levels with age and sex were determined using standard linear regression of inverse rank normalised protein levels. Analyses were adjusted for Olink analysis plate and sample age (time between biopsy and Olink assessment).

### pQTL analysis

Genome-wide association studies were performed for each protein in each tissue using REGENIE (16). We included the covariates age, sex, 10 genetic principal components (PCs, derived using FlashPCA (17)), Olink analysis plate and sample age (time between sample collection and proteomics analysis). In a sensitivity analysis for all tissues except mammary artery (where we only had CAD cases), we adjusted for CAD status.

### Significance threshold for cis-pQTLs

The p-value threshold representing empirical 1% alpha threshold for *cis-*pQTLs across all proteins (*i.e.* the total experiment) was calculated by permuting a normally distributed phenotype (to represent protein levels) and running an association test of this phenotype with the STARNET genotype data for the cis region (+/-500kb) of each gene that encodes the corresponding protein. This was repeated 1,000,000 times and the p-value threshold at which 1% of “proteins” had a significant genetic association meeting this threshold was identified (7×10^−6^). This was the same p-value threshold as the p-value threshold corresponding to a 1% false discovery rate when combining all results from the cis gene region (+/-500kb) of all proteins (including all SNPs from overlapping cis regions). Independent signals were identified using LD-based clumping in Plink (18), using a UKB reference panel. The r^2^ cut-off was 0.01.

### PheWAS, Colocalization and Mendelian randomization

To identify 1. shared genetic regulation of protein levels across tissues, 2. shared genetic regulation between gene expression and protein level in the same tissue, and 3. tissue specific protein contributions to cardiometabolic disease, we performed colocalization with cardiometabolic traits using Hypothesis Prioritisation for multi-trait Colocalization (HyPrColoc) and used a posterior probability of ≥0.7 to define colocalization (P_A_ and P_R_ were set at 0.7) (19). For analysis of shared regulation of proteins across tissues, we included all tissues in the same model for any protein with a *cis*-pQTL in at least one tissue. Pairwise colocalization of protein with eQTLs from the same tissue was performed using coloc and data from GTEx (20) and the published STARNET eQTL analysis of CAD cases (11) to identify shared regulation of gene expression and protein in the same tissue. We tested for colocalization with CAD-related biomarkers LDL, HDL, triglycerides and CRP using PanUKB data (21). For Coronary Artery Disease, we used the largest GWAS from a European population (22). For the colocalization with cardiometabolic traits, we included all tissues in the model (HyPrColoc, using the same parameters as for protein levels across tissues) and analysed any protein with a pQTL in at least one non-plasma tissue. To identify novel tissue-specific protein-trait associations, we performed PheWAS across all traits from the panUKB initiative (21), for the top significant independent SNPs for each protein where we had a pQTL in tissue but not in plasma. We then performed colocalization of all traits identified with protein levels in the specific tissue using HyPrColoc.

Mendelian randomization was performed using the TwoSampleMR package (23). Independent instruments were identified using LD-based clumping using an r^2^ threshold of 0.01. Where more than one independent QTL was identified, the inverse variance weighted (IVW) method was employed and for one independent SNP the Wald ratio estimate was calculated. Where more than two independent SNPs were identified, we performed weighted median, weighted mode and MR-Egger analyses to test for potential invalid instruments (24–26).

## Results

### Identification of tissue-shared and tissue-specific pQTLs

We analysed up to 284 individuals with plasma and/or tissue samples. Of these participants, 77% were male, with a mean age of 65 years (SD +/-11), and 70% had CAD at the time of surgery. Fourty-nine proteins in plasma were measured below the limit of detection (LoD) in more than 80% of individuals, compared to 38 in aorta, 28 in liver, 41 in mammary artery and 65 in muscle *(Supplementary Tables ST2-ST6).* This is much lower than the 382 proteins measured below LoD in more than 80% of individuals in the UKB-PPP (7), which likely reflects the high quality, and immediate preservation, of blood and tissue in STARNET. In total, five proteins were measured below LoD in more than 80% of individuals in all tissues. Using <80% missingness to determine presence of a protein in a tissue, 39 proteins were detected in at least one tissue but not plasma, and 16 proteins were detected in plasma only (*Supplementary Figure 1)*. Associations of plasma proteins with age and sex were comparable to those observed in UK Biobank pharma proteomics project (UKB-PPP), with 12 of the top 20 most strongly age-associated proteins in STARNET being in the top 20 most strongly age-associated in UK Biobank (*Supplementary Figure 2a).* Sex associations were even more consistent, with 16 of the top 20 being shared. SPINT3 and KLK3 levels were higher, and PZP lower, in males across all tissues (*Supplementary Figure 2b)*.

Due to less protein variability in tissues than in plasma *(Supplementary Table ST7)*, despite having similar sample sizes across the different tissues (276 for plasma, 272 for aorta, 262 for skeletal muscle, 241 for liver and 191 for mammary artery, with proteomic, covariate and genotype data), we detected a much higher number of plasma pQTLs than tissue pQTLs. Specifically, we identified more than three times the number of pQTLs in plasma, compared to liver, which had the next highest number (*Figure 1*). None of the proteins with pQTLs were those detected below LOD in >80% individuals.

**Figure 1:**
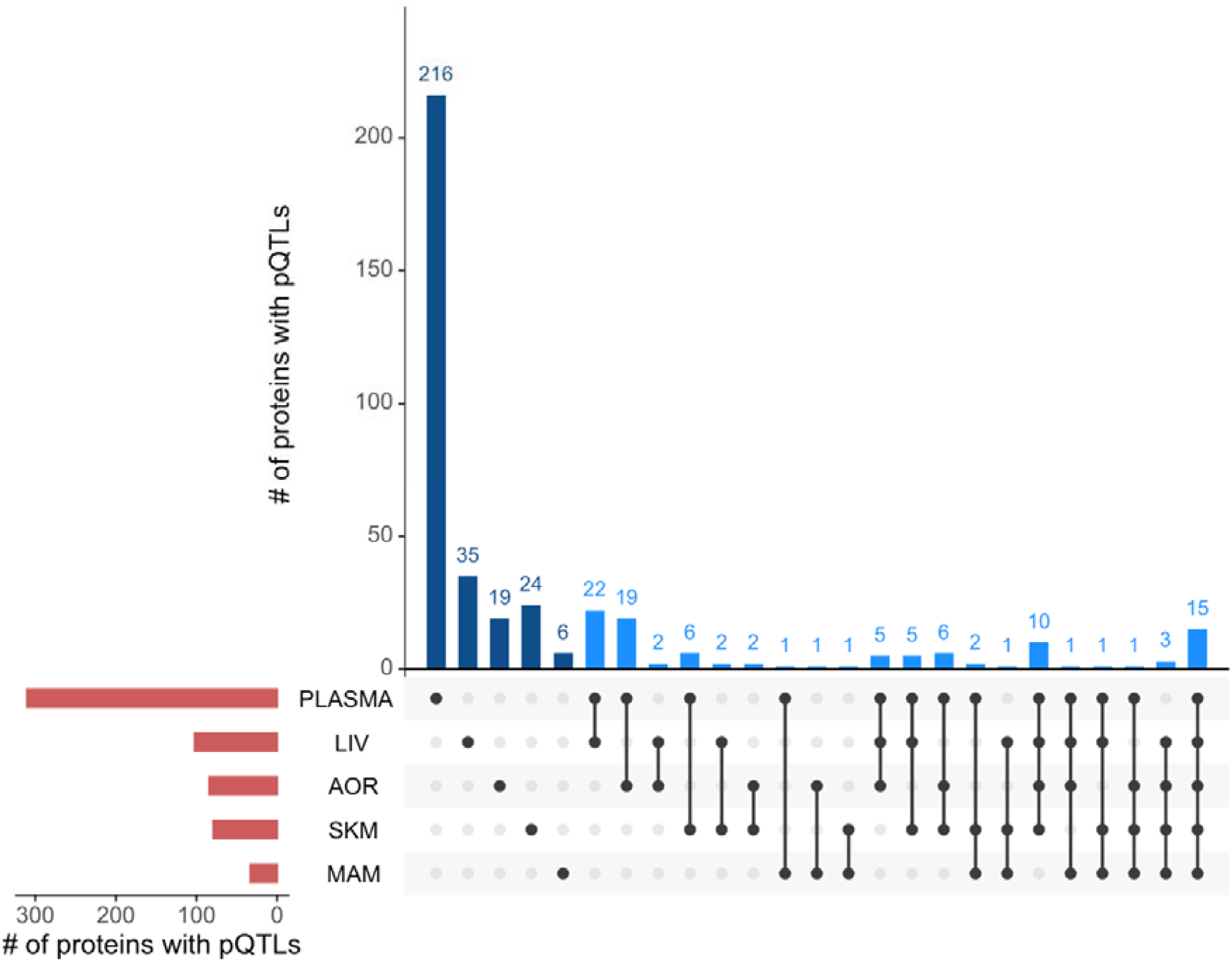
Distribution of proteins with cis-pQTLs across tissues LIV: liver, AOR: aorta, SKM: skeletal muscle, MAM: mammary artery

Of the 310 proteins with a *cis*-pQTL in plasma, 94 also had *cis*-pQTLs in other tissues, with the highest overlap observed in the liver, where 59 proteins had pQTLs in both liver and plasma. Fifteen proteins had a *cis*-pQTL across all tissues analysed. In total, 190 proteins had pQTLs in non-plasma tissues, and of these, 96 proteins did not have a pQTL in plasma in this analysis. On comparison with the UKB-PPP data, 13 of these 190 did not have *cis-*pQTLs in UKB (based on the significance threshold presented in the paper). Despite our study population being enriched for CAD, adjustment for CAD status made little difference to the results (*Supplementary Figure 3).* At an FDR threshold of 0.01, we identify 763 *cis*-pQTLs in total (*Supplementary Figure 4)*.

After performing LD-based clumping, the median number of independent genetic associations for each tissue was one. For aorta, 15 of the 84 proteins with *cis*-pQTLs had more than one independent *cis*-pQTL (p≤ 7×10^−6^), with four proteins having three independent genetic associations *(Supplementary Table ST8)*. In liver, 12 of the 102 proteins had more than one independent SNP, with three having three independent SNPs *(Supplementary Table ST9)*. In muscle, nine out of 79 had more than one independent genetic association, while ABO had four independent associations *(Supplementary Table ST10)*. Seven proteins for mammary had two independent SNPs with a p-value less than 7×10^−6^ *(Supplementary Table ST11)*.

### Tissue pQTLs were generally consistent in direction with plasma pQTLs

Out of 190 proteins with a pQTL in at least one non-plasma tissue, 106 did not share the same variant across different tissues (*Figure 2*), suggesting these proteins are influenced by distinct genetic factors. Most proteins that did share a causal variant across tissues were influenced by pleiotropic variants that affect multiple tissues at once. Specifically, 21 proteins shared variants across all tissues, including plasma, while four proteins had shared pQTLs in all non-plasma tissues. The liver shared variants with plasma for 48 proteins; the aorta and plasma for 44 proteins; skeletal muscle and plasma for 38 proteins; and mammary tissue and plasma for 31 proteins.

**Figure 2:**
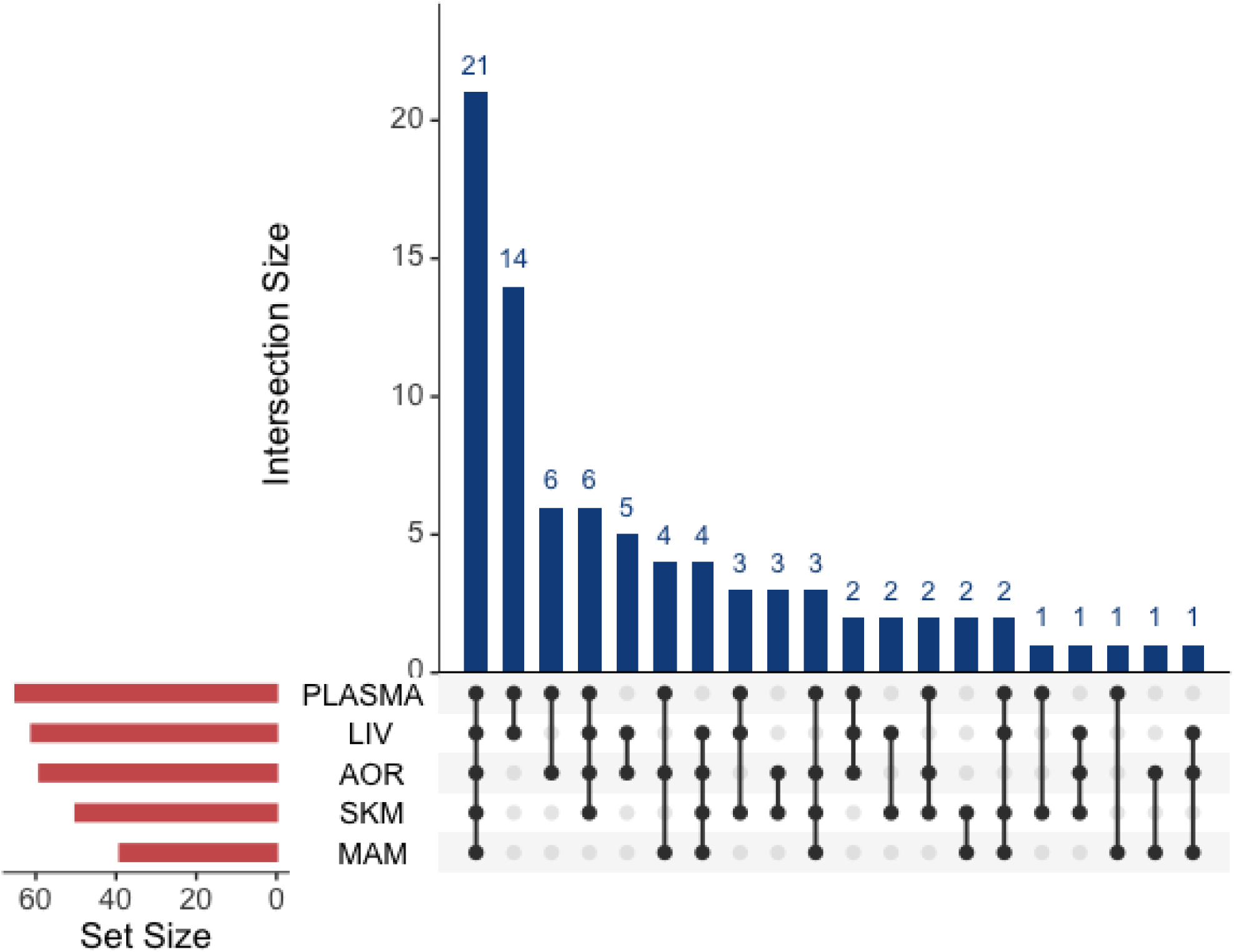
Upset plot showing number of proteins that shared causal variants (colocalized) across tissues Abbreviations: MAM: mammary artery, SKM: skeletal muscle, AOR: aorta, LIV: liver Set size= number of proteins with at least one pQTL. Intersection size= number of proteins with any pQTL in the relevant tissues (but not necessarily the same pQTL)

Generally, the directions of effect were the same in different tissues and in UKB plasma samples *(Figure 3)*, which was also true for plasma from the STARNET population (*Supplementary Figure 5*). When pQTLs were shared across tissues, the directions and magnitudes of their effects were consistent. However, there were a few exceptions. Exceptions, where the same variant was shared but had opposing effects, include CPPED1 in aorta, CA5A in liver, S100A16 in liver, and DUSP13 in skeletal muscle *(Figure 3)*. Other cases where sentinel SNPs in tissue had an opposing direction of effect in plasma reflected different genetic variants influencing plasma and tissue proteins (i.e. lack of colocalization) *(Supplementary Figure 6*). All sentinel SNPs for proteins with pQTLs in the mammary artery showed the same direction of effect in plasma *(Figure 3, Supplementary Figure 5)*.

**Figure 3:**
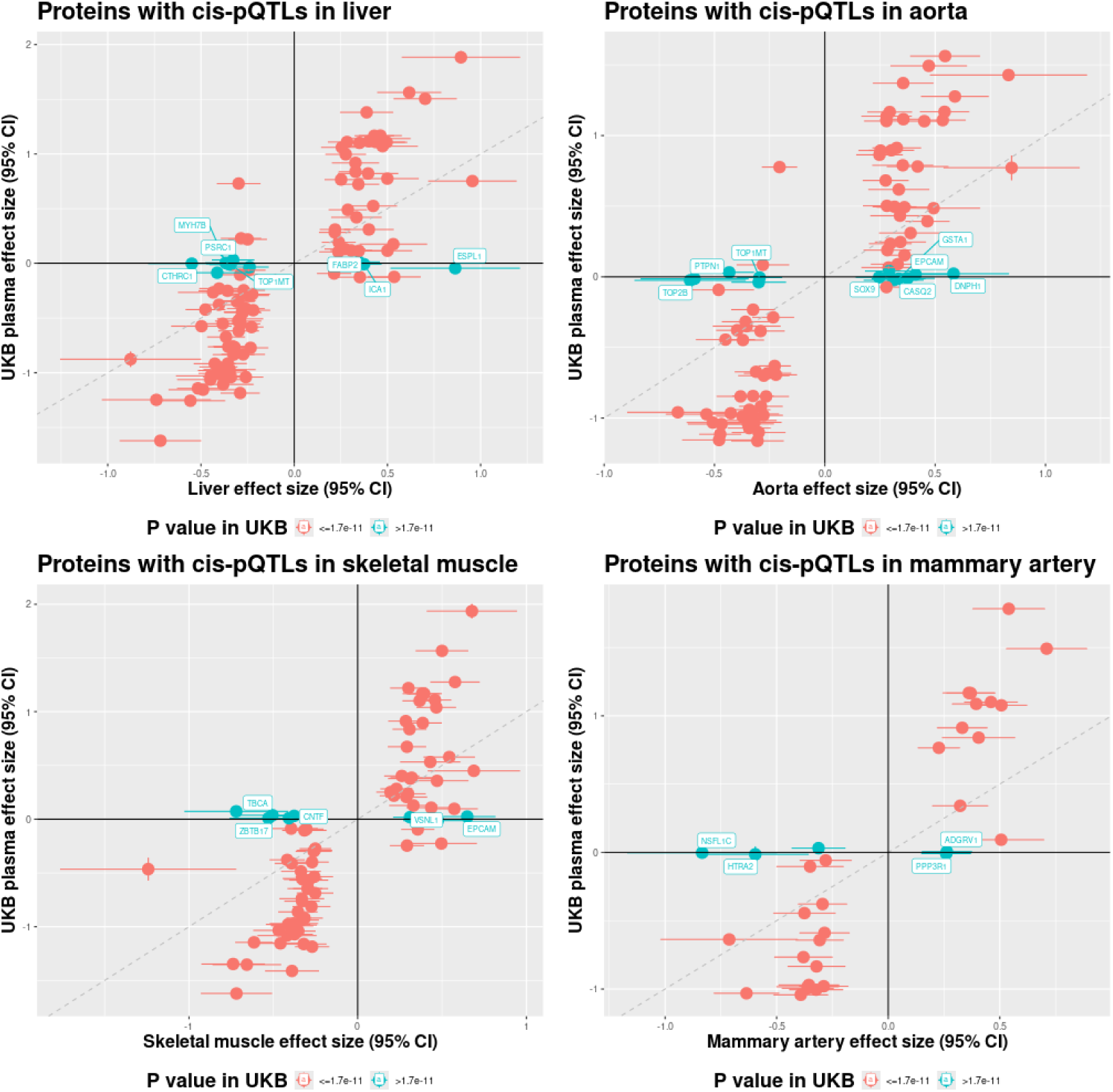
Comparison of effect sizes between STARNET tissues (x-axis) and UKB plasma (y-axis) for all proteins with cis-pQTLs in the tissue represented. The points represent the SNP with the lowest p-value in the cis region for the tissue. Labelled points represent proteins where there is a cis-pQTL in tissue but the same SNP does not reach p<0.05 nominal significance threshold in plasma in UKB.

### A large proportion of tissue pQTLs are not eQTLs in the same tissue

Of the 84 proteins with cis-pQTLs in aorta in STARNET, expression of 11 of the genes encoding them were not detected in aorta in either GTEx or STARNET (non-overlapping sample), as evidenced by the lack of eQTL data (*Supplementary Table ST8*). Twenty-two proteins colocalized with expression in GTEx and/or STARNET at PP≥0.7. Twenty-seven proteins had an eQTL at p≤7×10^−6^ but did not colocalize, and 24 proteins did not have an eQTL in GTEx or STARNET. Of the 102 liver proteins with cis-pQTLs, nine were not detected at the mRNA level in GTEx or STARNET *(Supplementary Table ST9)*. Thirty-two colocalized, 24 had a cis eQTL but didn’t colocalize and 37 did not have a liver eQTL. Of the 79 muscle proteins, 15 weren’t detected at mRNA level in GTEx or STARNET, 22 colocalized, 28 had an eQTL but didn’t colocalize, and 14 did not have an eQTL *(Supplementary Table ST11)*.

Of the pQTLs and eQTLs colocalizing in the same tissue, we observed several examples where the sentinel pQTL had opposing directions of effect on mRNA abundance versus protein levels *(Supplementary Figure 7)*. These include DPP7 in aorta, MIF and CA5A in liver and CNTF, DUSP13 and HDGF in muscle. The sentinel SNP for DPP7 (rs10747049) is a missense variant, associated with decreased protein in both aorta and plasma (7) and increased expression in aorta (20) and blood (27). This variant is predicted as likely benign by alpha-missense (28). There is additional colocalization with a splice QTL in the same tissue (29). The HDGF sentinel SNP is in complete LD with the missense variant rs4399146; this variant is also predicted to be likely benign (28). The DUSP13 sentinel SNP, rs6480771, is also a missense variant that is predicted to have a benign effect on protein function (30). The CA5A sentinel SNP is in high LD with the synonymous variant rs7186698 (r^2^=0.98). Although this variant does not result in any functional alteration to the protein, the codon is altered to a less frequent codon (31), which could result in a change in protein levels via differences in tRNA frequency (32), gene expression could therefore be upregulated as a compensatory mechanism. The opposing direction of effect of the sentinel SNP on protein levels versus gene expression for MIF in liver was also identified in an analysis of MIF protein in GTEx liver samples (33).

### A small proportion of tissue pQTLs are tissue specific

We identified 13 proteins with tissue-specific pQTLs (*i.e.* pQTLs in a non-plasma tissue) that had no pQTL in plasma in STARNET nor reported significant by the UKB-PPP (7) (*Supplementary Table ST12*). Of these, 11 still did not have *cis-*pQTLs in UKB at the significance threshold we employed in our analyses, highlighting their tissue specificity. Among these tissue-specific pQTLs, four proteins (PNMA2, ICA1, ESPL1 and MYH7B) had pQTLs only in the liver, three (PTPN1, SOX9, CASQ2) only in the aorta, three (PPP3R1, ADGRV1, HTRA2) only in the mammary artery, one (CNTF) only in skeletal muscle and one protein (EPCAM) had pQTLs in aorta and skeletal muscle only, and one (TOP1MT) in aorta and liver only. Eight of the 13 proteins had a cis-eQTL in the corresponding tissue in GTEx or STARNET, and four colocalized (SNP with p≤7×10^−6^, *Supplementary Figure 8*). Forty-eight proteins had a pQTL in only one of the non-plasma tissues investigated, in addition to a plasma pQTL. Of these, 28 colocalized with plasma (*Supplementary Figure 9*).

Of the 13 proteins with tissue-specific pQTLs, we identified examples where the pQTLs map to tissue-specific regulatory regions (*Supplementary Figure 10).* For example, one of the sentinel SNPs, rs139120998, for ESPL1 in liver specifically, is in a regulatory element specific to liver. Additionally, the variant rs17568220, which is in high LD (r^2^ 0.99) with the sentinel SNP for CNTF in muscle, is in a muscle-specific regulatory region. Equally, of the proteins with pQTLs only in one tissue and plasma, CCL16 and IL17RB have pQTLs in regulatory regions specific to liver, highlighting a potential tissue-specific contribution to circulation.

### Identifying Tissue-Specific Effects on Cardiometabolic Traits

To identify tissue specific effects on cardiometabolic traits, we performed colocalization of significant pQTLs with CAD, CAD-related biomarkers and BMI to identify shared genetic drivers of proteins in tissue and cardiometabolic disease. PNLIPRP2 protein levels in all tissues colocalized with LDL levels, CTSH in both liver and plasma colocalized with LDL whereas PKN3, PRSS53, PSRC1 and SORT1 colocalized with LDL but only in the liver (*Supplementary Figure 11*). SORT1 and PSRC1 in liver also colocalized with CAD, HDL, CRP and triglycerides *(Figure 4, Supplementary Figure 11)*. MIF protein levels in liver only and GSTT2B in all tissues colocalized with both triglycerides and CRP. Other cross-tissue colocalizations were seen for CD300LF and CRP, CFHR2 and HDL, MEP1B (excluding liver) and HDL and LILRB5 and triglycerides. Skeletal muscle specific colocalizations were observed for CFHR4 and HDL, and AOC1 and HDL. Aorta specific colocalizations were observed for MET and triglycerides and CTRL and triglycerides. Other examples of shared genetic drivers were between protein levels in one tissue and plasma and a cardiometabolic outcome, included LILRB2 in aorta and both HDL and triglycerides, CDA in liver and HDL and GSTA3 in liver and triglycerides. None of the proteins with a pQTL in a non-plasma tissue colocalized with BMI.

**Figure 4:**
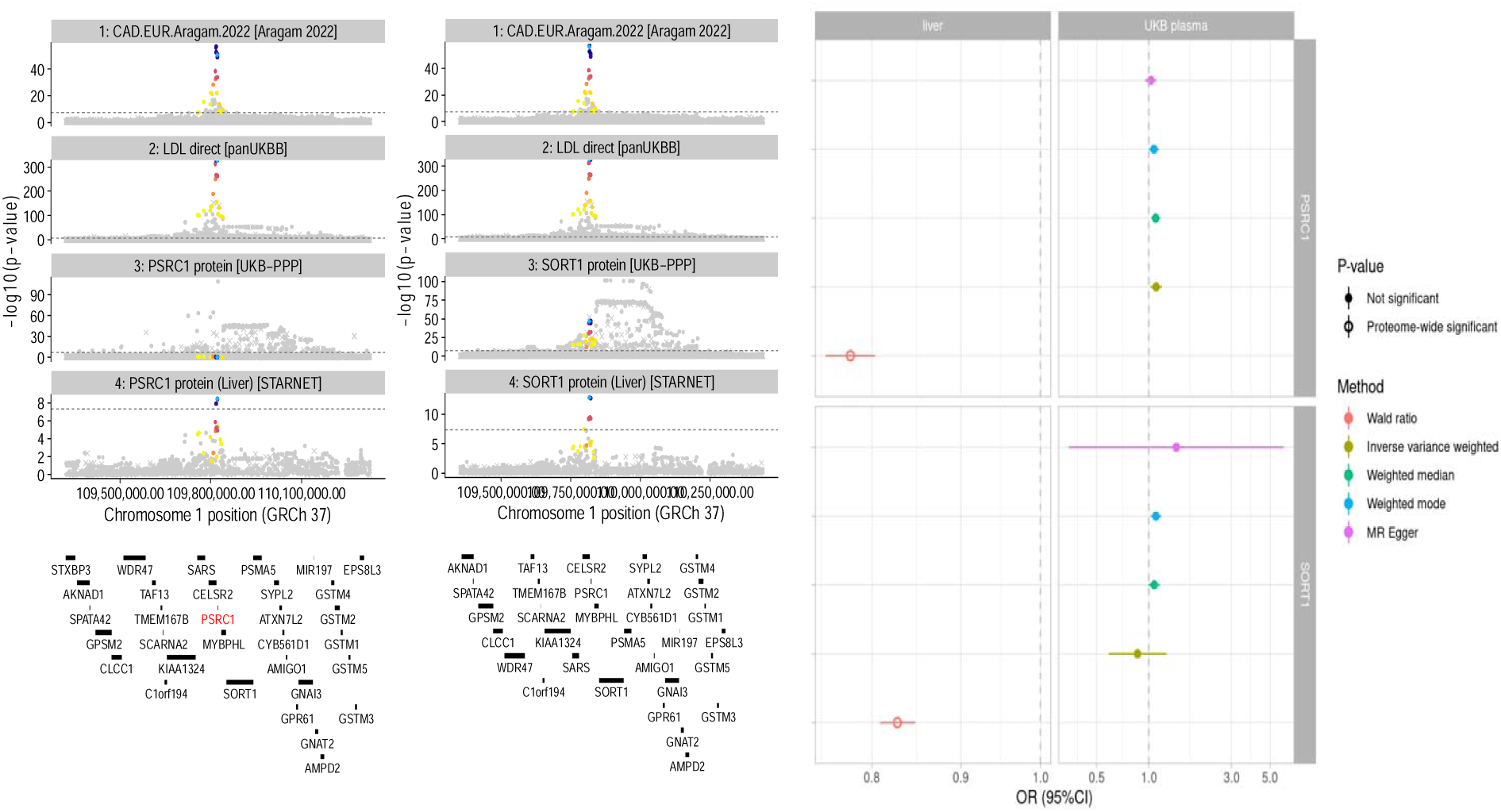
Regional association plot showing the sharing of genetic association between CAD and PSRC1 and SORT1 protein levels in the liver specifically and the Mendelian randomization effects of blood vs liver proteins on CAD risk

Mendelian randomization analyses showed a liver-specific effect of increasing SORT1 and PSRC1 protein on lower risk of CAD (*Figure 4),* as well as increased HDL and CRP levels and reduced plasma triglycerides *(Supplementary Figure 12)*. The effect of increasing PKN3 in liver on lower LDL was also seen for plasma in UK Biobank *(Supplementary Figure 12)*. Increasing genetically predicted PRSS53, in both liver and plasma, and PNLIPRP2, in all tissues, also putatively increases LDL. For HDL, there were opposing effect estimates for skeletal muscle vs plasma, with increasing AOC1 in skeletal muscle suggested to decrease HDL whereas increasing levels in plasma putatively increase HDL in circulation *(Supplementary Figure 12)*. There was also evidence for an effect of increasing CFHR2 in liver, muscle and plasma on decreased HDL, whereas the effect of CFHR4 was inconsistent across tissues. Increasing LILRB2 in aorta and plasma putatively decreases circulating HDL, as does increasing MEP1B in mammary artery, muscle and plasma *(Supplementary Figure 12)*. The opposite direction of effect was observed for LILRB2 and triglycerides. Increasing GSTA3 and LILRB5 in liver putatively decreases plasma triglycerides, whereas increasing GSTT2B and MIF increases triglycerides. These effects were seen in at least one other tissue. Increasing CTRL in aorta and plasma putatively decreases triglycerides according to our MR analyses. Effects of MIF and GSTT2B in liver on plasma CRP were also observed, although the direction of effect opposed triglycerides. Finally, we also observed an effect of increasing CD300LF in muscle and plasma on increased levels of plasma CRP.

PheWAS of the independent pQTLs identified in tissues, for proteins that lacked a pQTL in plasma, identified several associations in addition to those described above. Notably, the top SNP for MYH7B in liver was associated with traits such as IGF-1 and skin cancer *(Supplementary Figure 13)*. However, the lack of colocalization of the traits with MYH7B due to the LD structure in the region make it difficult to determine if variants causing change in MYH7B protein levels are causal for these traits (*Supplementary Figure 14*). A proteome wide association analysis of these variants also highlighted some potential circulating biomarkers likely to be altered by changes of levels of these proteins in tissues, such as circulating NEFL as a biomarker of changing CNTF in muscle (*Supplementary Figure 15)*.

## Discussion

The plasma proteome holds great promise to better understand mechanistic underpinnings of complex diseases, and to define useful biomarkers. However, without better insights into the tissue-specific regulation of proteins being secreted or leaked into the circulation, our understanding of the plasma proteome will remain incomplete. To our knowledge, using the unique features of multi-tissue sampling from living patients in the STARNET cohort, our study is the first to identify multi-tissue, and tissue-specific pQTLs, providing new insights into the genetic regulation of protein levels in the arterial wall and metabolic tissues in relation to plasma proteins and cardiometabolic traits.

By uncovering tissue-specific pQTLs, we can identify and validate new drug targets that are not detectable in plasma. We identified novel tissue-specific pQTLs, which were not detected in plasma, even in a large, well-powered, analysis of the UK Biobank. This underscores the importance and utility of generating tissue-specific pQTLs to evaluate drug targets that are not secreted into blood. We identified more pQTLs in plasma compared to other tissues, which may be expected given the lower protein variability detected in the tissues compared to plasma. This likely reflects the fact that, unlike tissue where proteins are strictly regulated to fulfil certain organ functions, the plasma proteins vary along with the status of multiple organs throughout the body secreting or leaking proteins into the circulation (34). The greatest overlap in pQTLs, as determined by shared causal variants, was observed between liver and plasma. This finding aligns with liver being a major source of protein secretion into circulation (35). Additionally, we found that the majority of pQTLs in tissues were not eQTLs in the same tissue, consistent with a previous study, although in brain in relation to proteins in the cerebral spinal fluid (36).

We are aware of two previous studies that identified pQTLs at scale in tissue, however using mass spectrometry (MS) instead of affinity-based proteomics (33,37), which is more expensive and requires additional sample preparation. The He *et al* study, which focused on liver, included a similar number of individuals (287) and detected 1,344 proteins in >90% samples (37). In comparison, we identified 1,564 proteins with >90% measurements above the limit of detection in liver. Their analysis was performed genome-wide to identify both *cis-* and *trans-*pQTLs, and they identified *trans*-pQTLs for more proteins than they did *cis*-pQTLs (602 *vs* 74, respectively). We took the approach of testing fewer variants to maximise statistical power, which could explain the difference between the number of *cis*-pQTLs we identified. Only pQTLs reaching the genome-wide significance threshold were made available in the He *et al* study, preventing us from replicating or meta-analysing our associations. A recent preprint presented *cis*-pQTL findings from five non-plasma tissues (liver, colon, heart, lung and thyroid), with only liver overlapping this analysis. This paper used an FDR threshold of 0.01 to define a significant *cis*-pQTL, identifying 1,981 *cis*-pQTLs across five tissues (33). However, this threshold does not fully account for the number of statistical analyses performed as it does not account for the number of proteins measured. This study measured a total of 10,841 proteins via MS (in at least one tissue) (33), whereas we investigated 2,773 autosomal-encoded proteins. Similarly, a MS-based pQTL study has previously been performed for atrial appendage tissue in individuals with atrial fibrillation undergoing CABG surgery (38). This study detected 1,337 proteins in 75 individuals, with 45 *cis*-pQTLs (significant at a p<10^−5^ significance threshold) compared to 110 at this threshold in LIV in our study *(data not shown)*.

Plasma is a convenient biological sample for proteomics analysis due to its ease of collection and its composition as a pool of proteins secreted from, or released by, tissues (34). However, proteins that are not secreted from tissues, nor expressed by blood cells, are unlikely to be detected in plasma samples. For example, HMG-CoA reductase (encoded by the *HMGCR* gene) is the target of statins in the liver for lowering of cholesterol and prevention of coronary heart disease (39). As HMG-CoA reductase is not secreted into circulation, there are no pQTLs for this protein in plasma, despite its measurement in large, well-powered population studies using the Somalogic assay (40–42). Even where protein is present in both tissue and plasma, we highlight a few examples where estimates of direction of effect differ between tissue and plasma, with the minor allele associated with higher protein levels in tissue and lower levels in plasma, or vice versa. A potential explanation could be the function of these proteins; if they have roles in cell survival or protein secretion, it could explain why higher levels in the tissue would lead to lower levels secreted or leaked into plasma.

Incorporating tissue-specific proteomics in therapeutic target validation helps identify the tissue driving the effect on a disease outcome. For instance, the liver-specific pQTL for SORT1 underscores its potential as a therapeutic target for reducing coronary artery disease (CAD) risk through liver-specific mechanisms. The colocalization of the pQTL in liver with the eQTL in liver shows a mechanism by which the genetic variant acts on protein levels through gene expression, leading to a change in CAD risk. The role of the *SORT1-PSRC1* locus was discovered by early LDL (43,44) and CAD GWAS (45,46), with a key role of these genes in liver suggested by an association of the fine-mapped variants with *SORT1* and *PSRC1* expression in liver (47). The addition of our liver protein data substantiates a causal role of SORT1/PSRC1 protein in liver supporting it as a potential candidate drug target for CAD. This finding aligns with previous studies highlighting the role of SORT1 in hepatic lipoprotein production and CAD risk (47,48). Our data enable a more precise understanding of how genetic variations influence protein levels in specific tissues, paving the way for drugs to be designed to modulate protein levels in the target tissue, minimizing off-target effects and improving efficacy.

The addition of tissue-specific pQTL data can aid in determining the direction of effect a new drug should have on its target. To date, molecular QTL data have been used to estimate direction of effect using Mendelian randomization estimates of plasma protein and/or tissue gene expression, or an intermediate biomarker (49). However, we highlight several examples where MR would give opposing directions of effect between tissue gene expression and plasma protein levels. For example, higher MIF protein levels in liver appear to increase plasma triglycerides, whereas higher gene expression in liver putatively lowers triglycerides *(data not shown).* The direction of effect observed for protein is concordant with the effect observed in *in vivo* models (50). The direction of effect for CRP, however, was not consistent with the role of increased MIF in inflammation (51). There are several possible explanations for opposing directions of effect between gene expression and protein, including feedback loops whereby increased protein levels in tissue or blood could negatively regulate gene expression. Coding variants could result in reduced protein function or stability, leading to positive regulation of gene expression to counteract the reduced protein function or the reduced protein half-life. Coding variants that increase gene expression but have no effect on protein levels could equally be detected as pQTLs due to a slight change in structure that reduces antibody binding affinity. Tissue pQTLs are therefore an important layer of evidence when determining mechanism of action for targeting a particular protein.

Tissue-specific proteins and pQTLs can aid in the discovery of biomarkers that reflect changes in target tissues, facilitating the development of diagnostic tools and monitoring therapeutic responses (52). For example, the trans effect of CNTF protein in muscle on circulating NEFL levels highlights a potential biomarker for neuronal degeneration given that CNTF promotes neuronal survival (53) and NEFL is a marker of neuronal degeneration (54). This highlights the added advantage of studying *trans*-pQTLs for circulating proteins (when sample size provides sufficient statistical power).

Overall, our findings emphasize the need for tissue-specific proteomics in understanding the genetic basis of cardiometabolic disorders and developing more effective therapeutic strategies. By providing a comprehensive dataset of tissue-specific pQTLs, we hope to facilitate future research and clinical applications that will ultimately improve patient outcomes.

### Limitations

This study has some limitations. Our study identified a significant number of tissue-specific pQTLs; however, it is important to acknowledge the potential for false negatives due to our limited sample size. The relatively small cohort size may result in insufficient power to detect pQTLs for less abundant proteins or those with smaller effect sizes. This limitation is particularly pertinent in tissues where protein extraction and quantification may be more challenging compared to plasma. Due to the limited sample size, we also did not have statistical power to identify *trans-*pQTLs in this analysis *(Supplementary Figure 16)*. There is evidence to suggest that *trans-*pQTLs are more likely to be tissue-specific than *cis-*pQTLs (36,55). Importantly, the identification of biologically relevant pQTLs such as SORT1-PSRC1 provides support that our significance threshold for the identification of *cis*-pQTLs is appropriate. Although our population is enriched for CAD, the observed consistency of effect estimates when adjusting for CAD status and the general consistency in pQTL direction between diseased tissue such as the aorta and non-diseased tissues suggest that pQTL effects are not biased by CAD status.

Generation of pQTLs using proteins assessed by affinity-based techniques are vulnerable to potential epitope effects, whereby a genetic variant that alters amino acid sequence (*i.e.* a missense variant) changes the protein conformation at the site of one of the two antibodies binding, thus the genetic variant appears associated with protein levels when in fact the protein levels are not altered (41). These epitope effects could be tissue-specific, if the protein is only found in a specific tissue, or alternative splicing leads to removal of the protein altering variant in a tissue-specific manner. Of the tissue-specific pQTLs we identified (*i.e.* those without a pQTL in plasma) only the sentinel pQTLs for PNMA2 and MYH7B in liver were missense variants (30). SORT1-PSRC1 provides support for use of Olink for tissue proteomics, despite the development of the technology on plasma; the same sentinel SNP was identified in the GTEx MS pQTL analysis (33), validating the measurement of SORT1 by Olink in the liver. The sensitivity of Olink to detect proteins in tissues remains to be determined by studies performing MS and Olink analysis on the same tissues. Lastly, this analysis is based on a purely European sample which means we may not have identified some tissue-specific pQTLs if their frequency is rare in European populations.

## Conclusions

In summary, our study provides novel tissue-level protein expression data from cardiovascular disease patients and, combining with genetics, highlights the importance of considering tissue-specific protein regulation in therapeutic drug discovery and development. Our findings offer valuable insights when drug targets act through tissue-specific mechanisms and show how genetics can identify potential therapeutic target proteins when they are not detectable in plasma. To enhance future research, it may be beneficial to consider expanding sample sizes, integrating multi-omics data, longitudinal sampling in the same individuals, and these same analyses in diverse populations. These efforts will enhance the detection and validation of pQTLs, which will contribute to more effective and targeted therapeutic strategies, ultimately contributing to better clinical outcomes for patients.

## Author contributions

AEH: Conceptualization, Formal Analysis, Writing-Original Draft, KS: Tissue preparation, Methodology, Data Curation, Writing-Original Draft, SH: Methodology, Formal Analysis, MT: Conceptualization, Writing-Review and Editing, MGR: Conceptualization, Formal Analysis, Writing-Review and Editing, KEH: Methodology, Formal Analysis, HT: Validation, Writing-Review and Editing, AR: STARNET co-PI, Tissue sampling, Methodology, Data Curation, Writing-Review and Editing, EB: STARNET author, JLMB: STARNET PI, Conceptualization, Data Curation, Supervision, Writing-Original Draft, JMMH: Conceptualization, Funding acquisition, Supervision, Writing-Original Draft, YJ: Supervision, Writing-Original Draft

## Acknowledgements

With thanks to the STARNET and UK Biobank participants for providing their samples for inclusion in this analysis. Thanks to Reka Nagy for developing the tools used to create the regional association plots.

## Data availability

The raw proteomics data are not publicly available due to research participant privacy/consent. Summary statistics for all proteins in all tissues will be deposited in GWASCatalog at the time of publication.

