## Supplementary Figure for "Multi-Tissue Profiling Reveals tissue-specific protein regulation and relationships Between Protein Quantitative Trait Loci (pQTLs) and Cardiometabolic Disease"

**Supplementary Data**

*Supplementary Figure 1: Upset plot of the number of proteins with at least 20% of individuals having values above the limit of detection, across tissues*

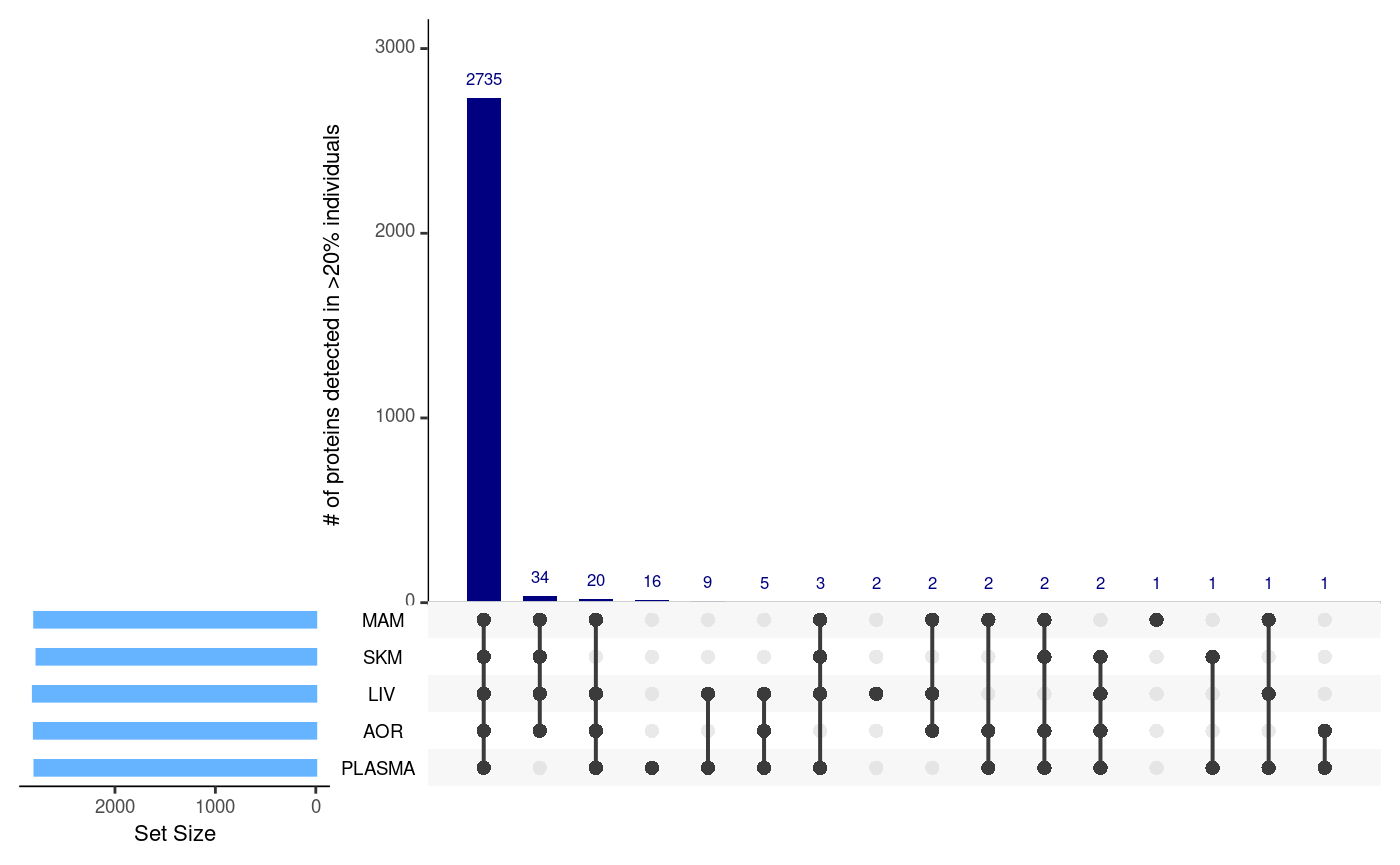

*Supplementary Figure 2: (a) Associations of plasma proteins with age and sex and comparison with associations in UKB-PPP and (b) associations of proteins with age and sex across tissues*

*(a)*

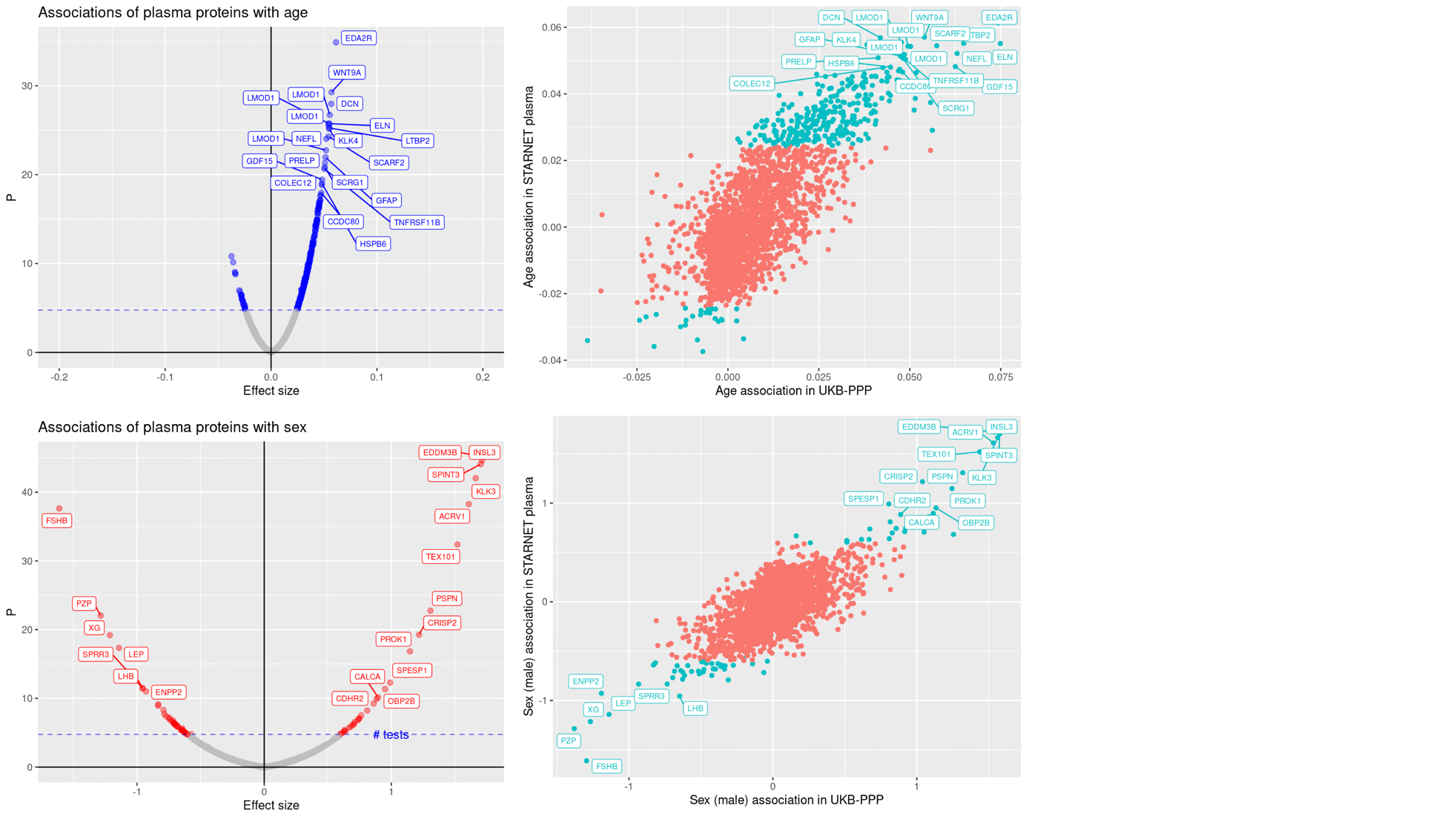

*(b)*

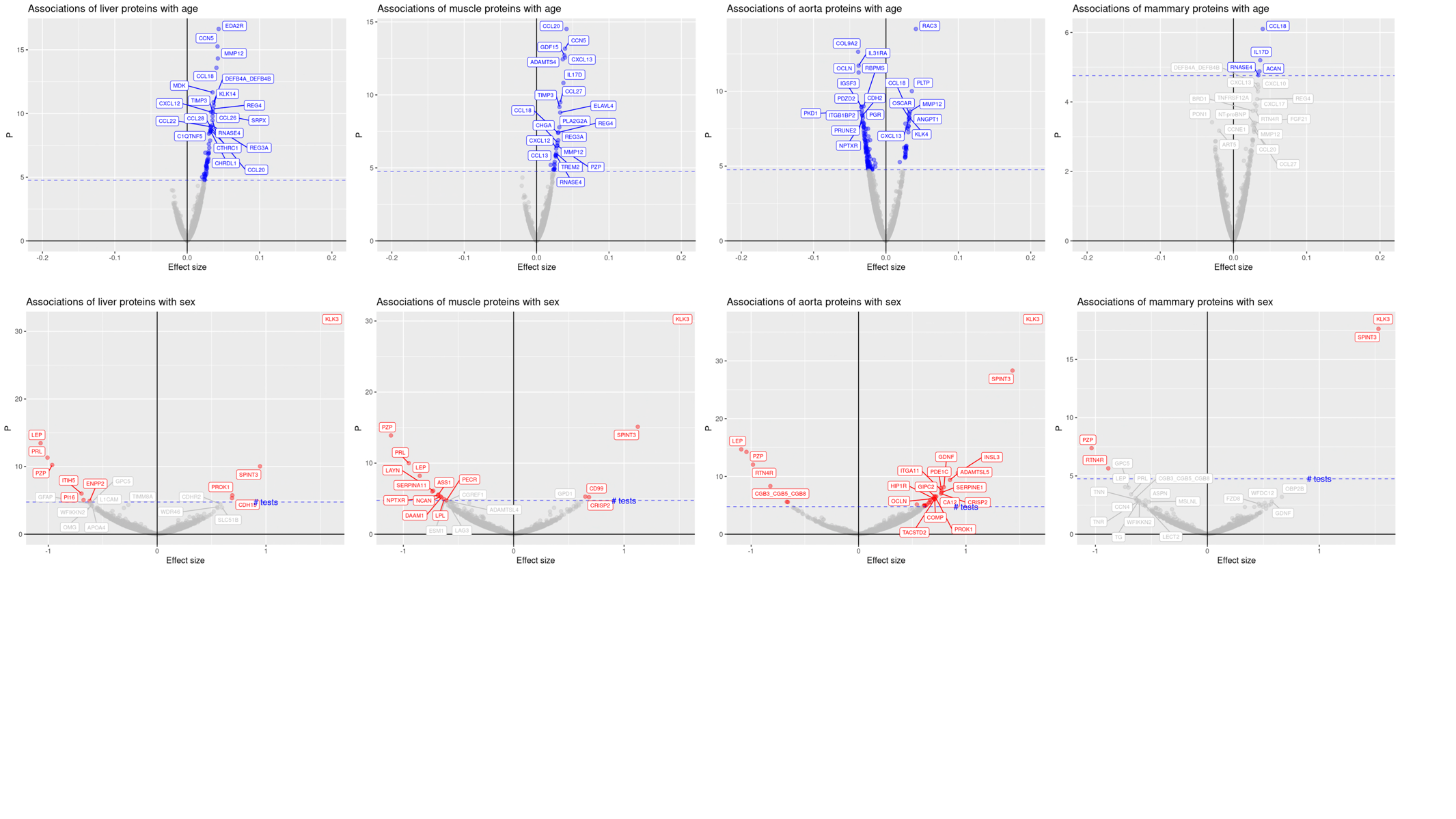

*Coloured points represent associations with a p-value less than 0.05/number of proteins analysed. The 20 proteins with the lowest p-values are labelled.*

*Supplementary Figure 3: comparison of associations of sentinel SNPs with protein levels before and after adjustment for CAD status*

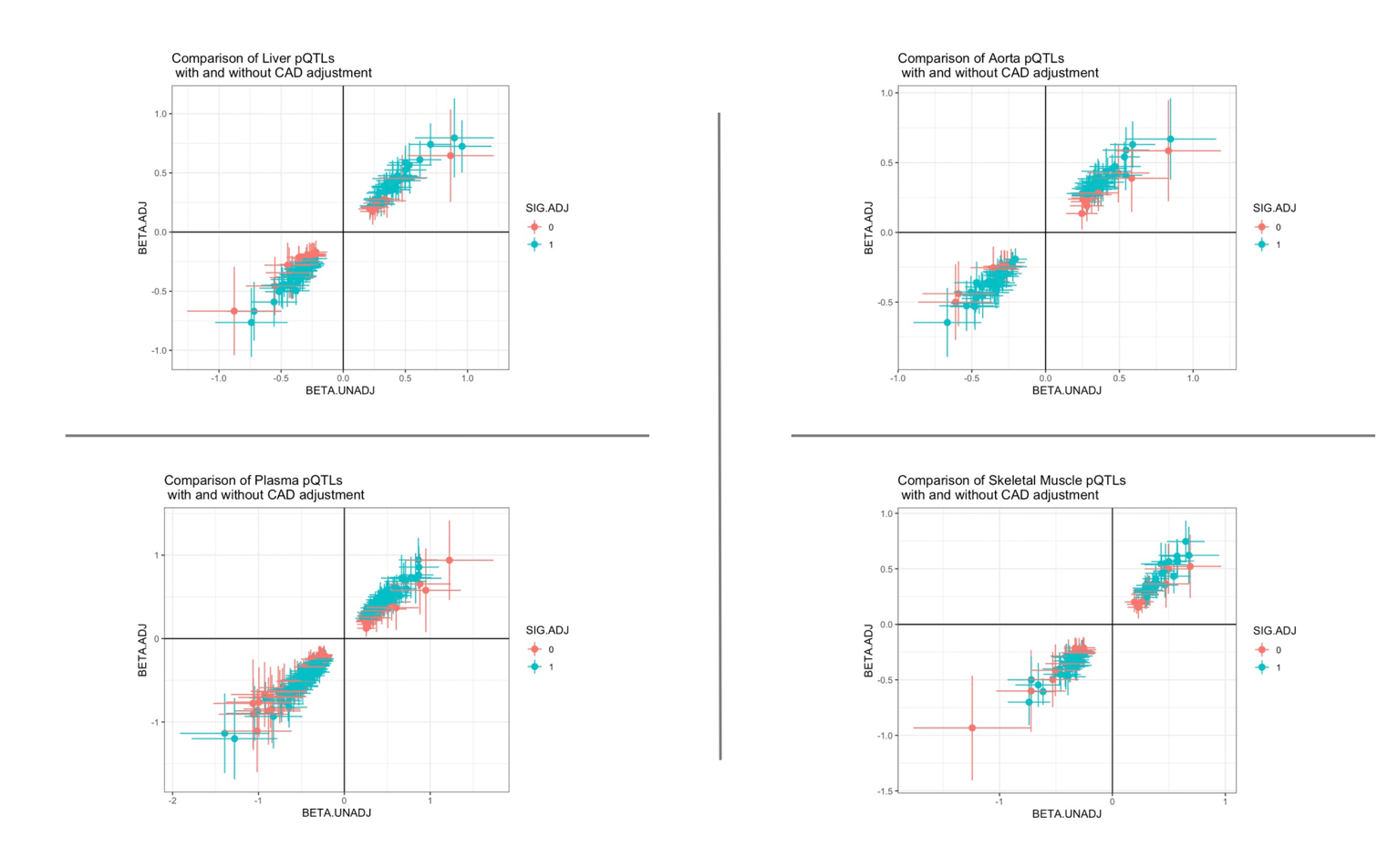

*Sig.Adj represents a p-value less than 7x10^-6^ in the CAD-adjusted analysis*

*Supplementary Figure 4: Upset plot of the number of cis-pQTLs identified using a false discovery rate threshold of 0.01*

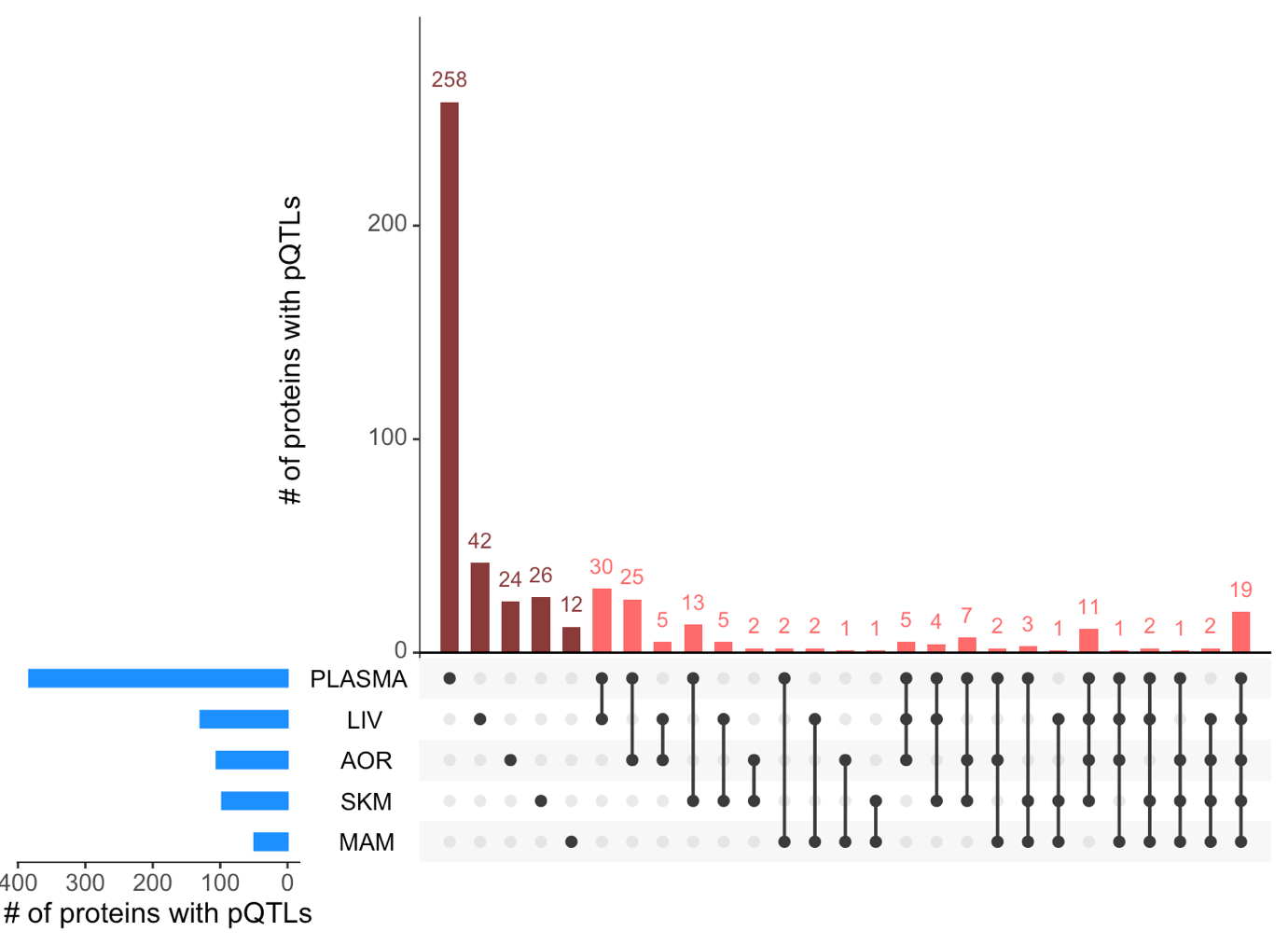

*Abbreviations: LIV: liver, AOR: aorta, SKM: skeletal muscle, MAM: mammary artery*

*Supplementary Figure 5: Comparison of effect sizes between STARNET tissues for proteins with a cis-pQTL in the relevant tissue. Points represent the SNP with the lowest p-value in the cis-region for the tissue of interest. Labelled points represent proteins where the SNP reaches the significance threshold in both tissues but has opposing directions of effect.*

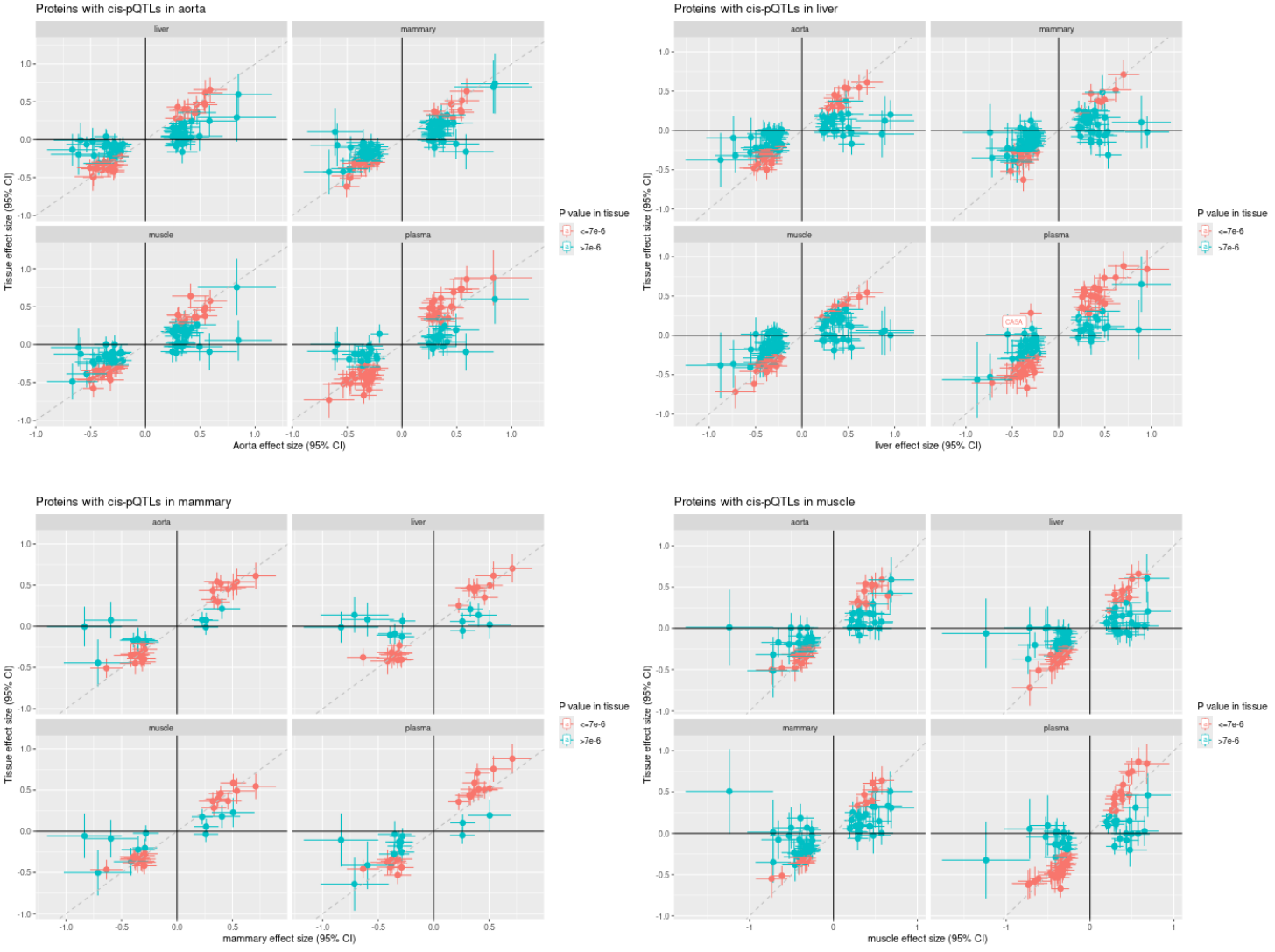

*Supplementary Figure 6: Regional association plots for proteins where the sentinel tissue SNP had opposing direction of effect on protein levels in plasma*

 
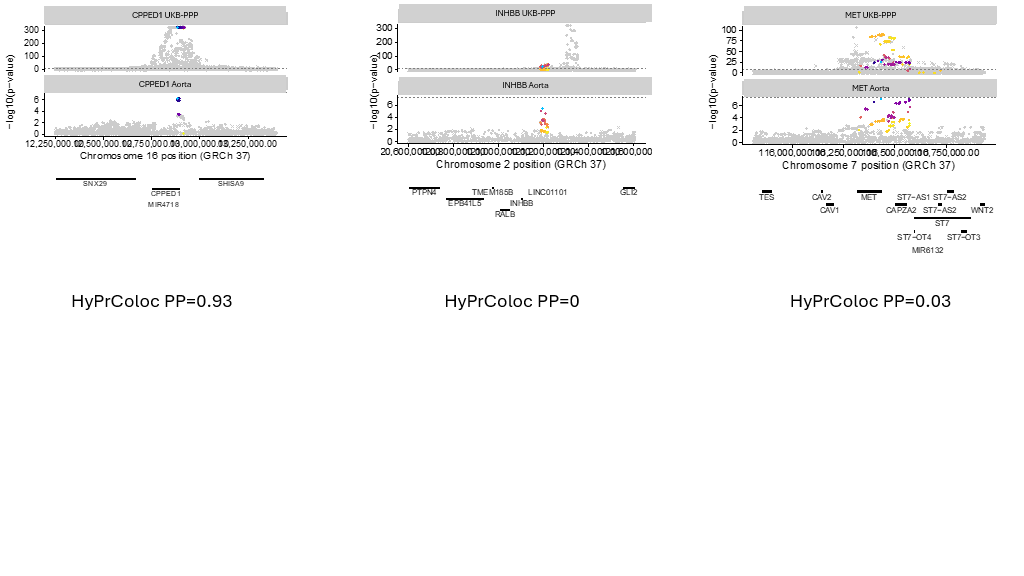
  
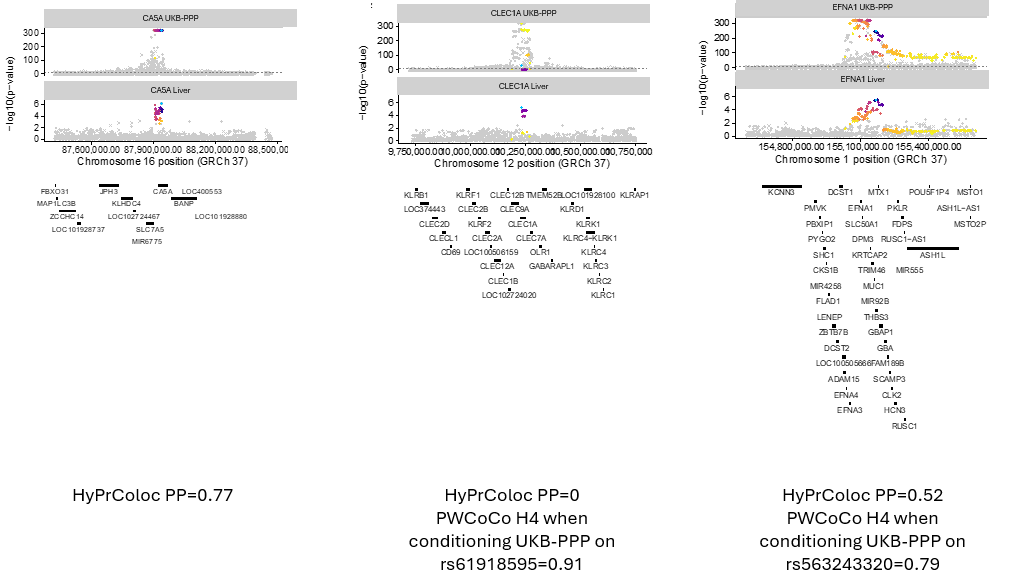
  
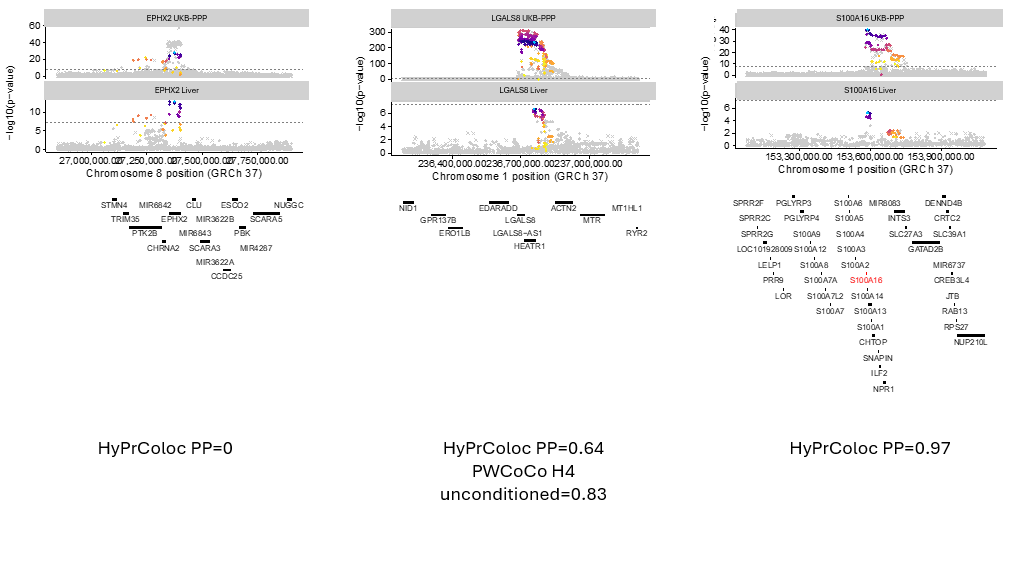
  
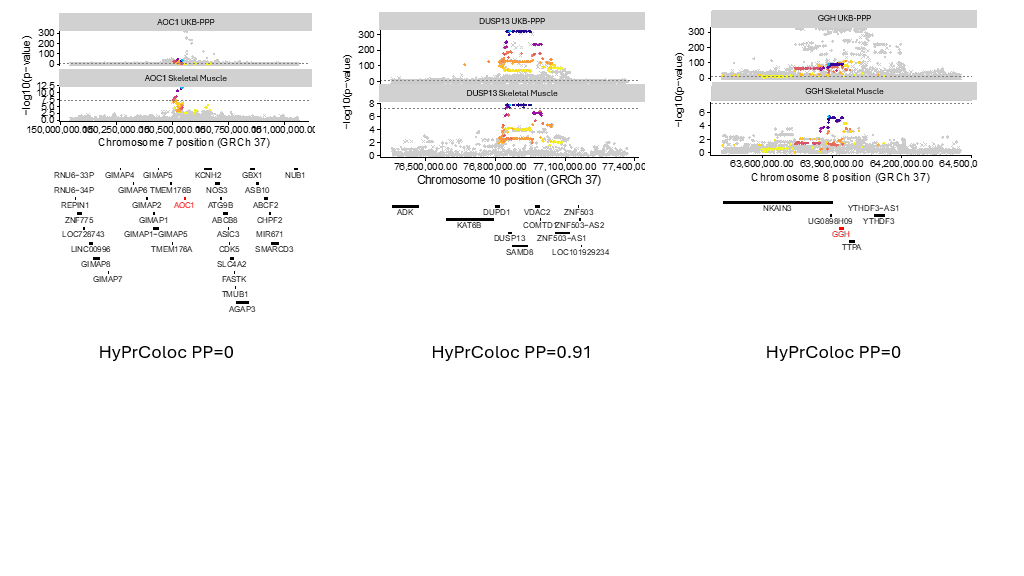

*Abbreviations: PP=Posterior Probability*

*Supplementary Figure 7: Forest plots for the sentinel pQTL in tissue showing opposing directions of effect on gene expression versus protein levels*

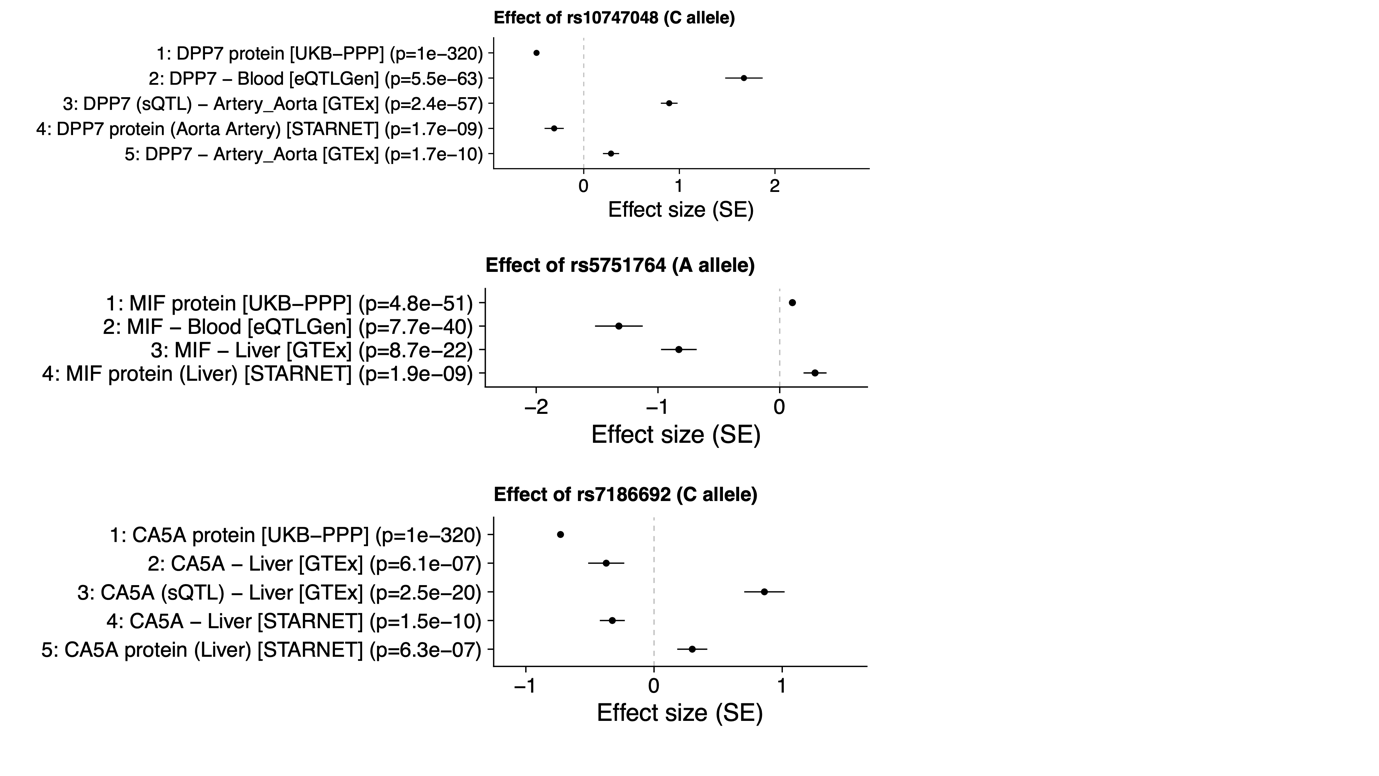

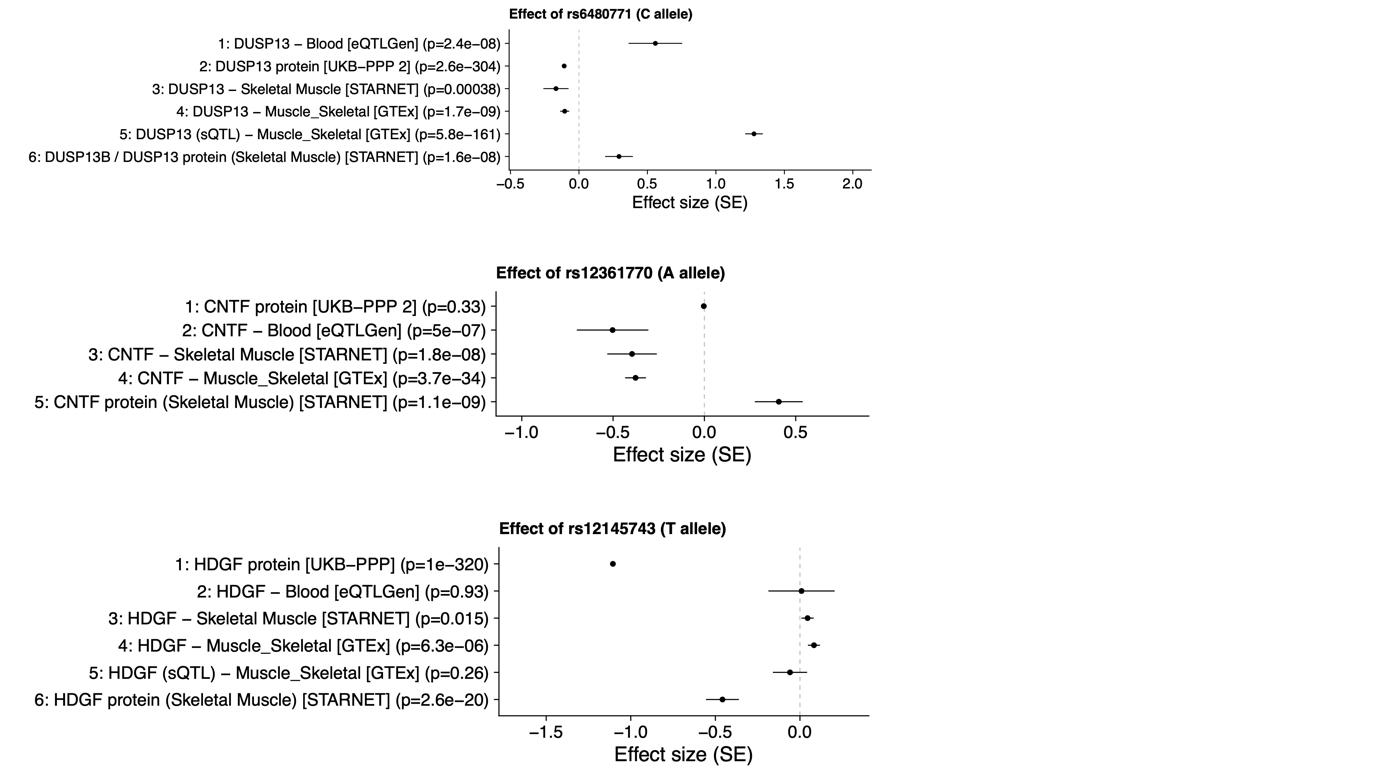

*Abbreviations: sQTL: splicing quantitative trait locus*

*Supplementary Figure 8: Regional association plots for proteins, and gene expression of the encoding gene, with a cis-pQTL in tissue but not plasma*

*Supplementary Figure 9: Regional association plots for the proteins with pQTLs in plasma and one other tissue*
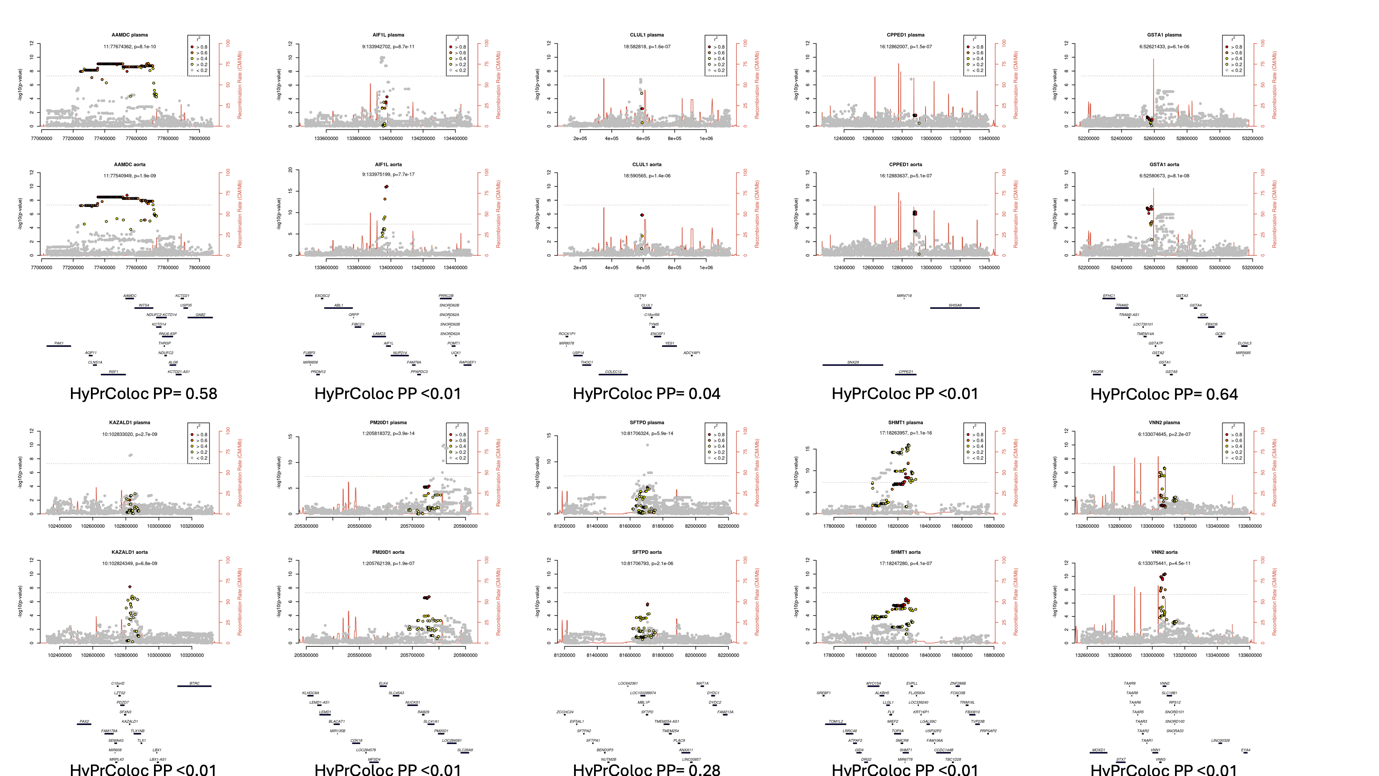

 
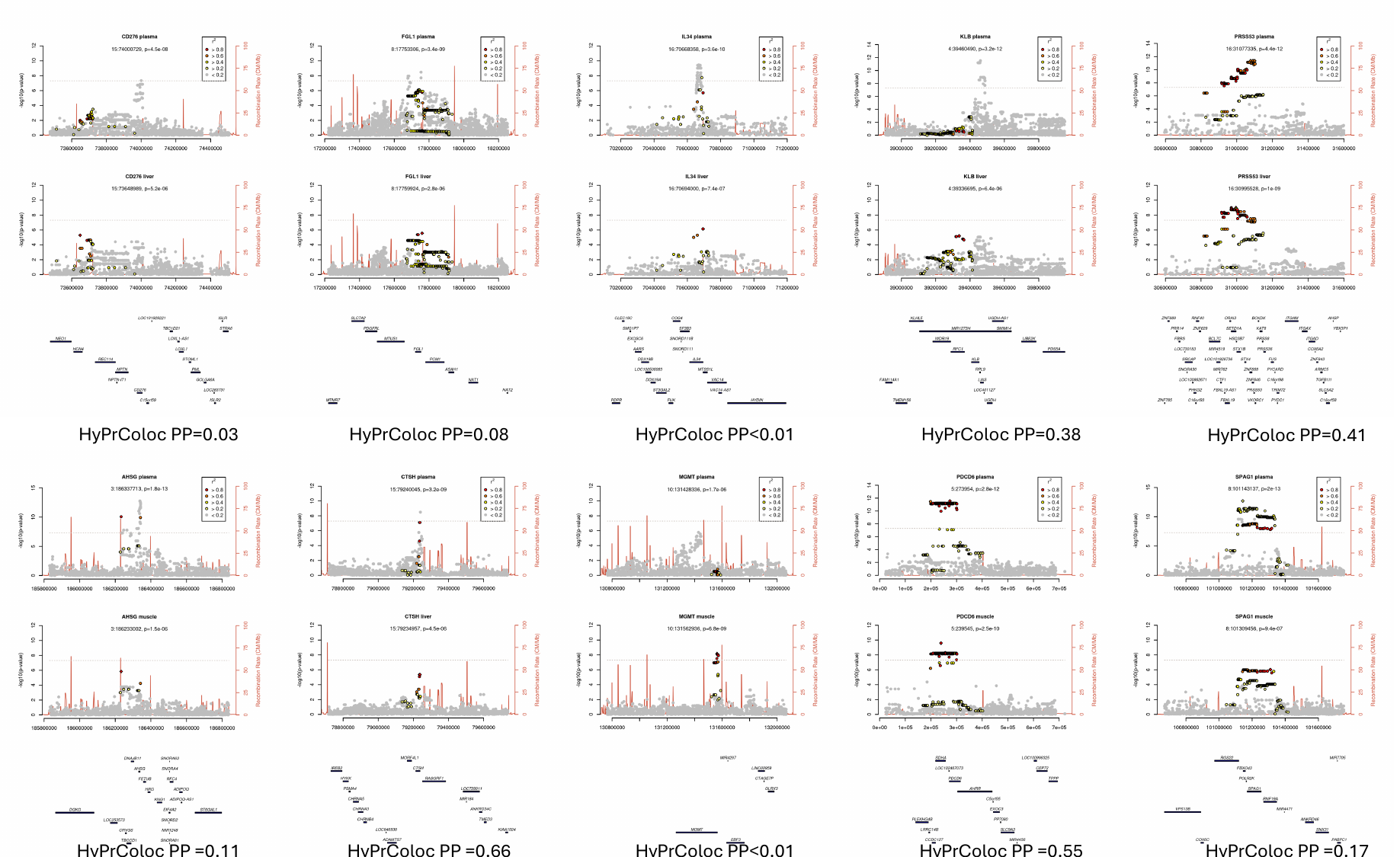

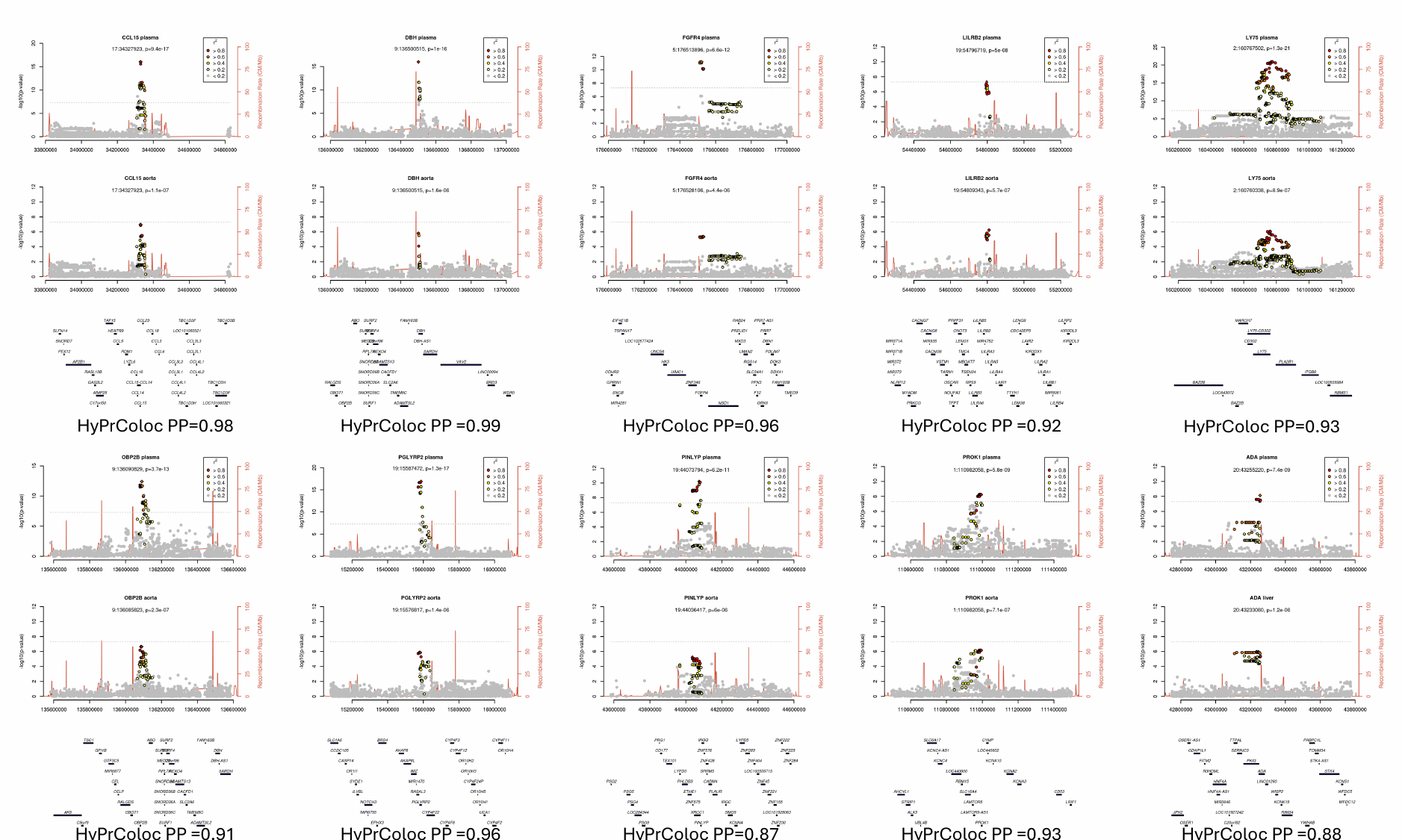

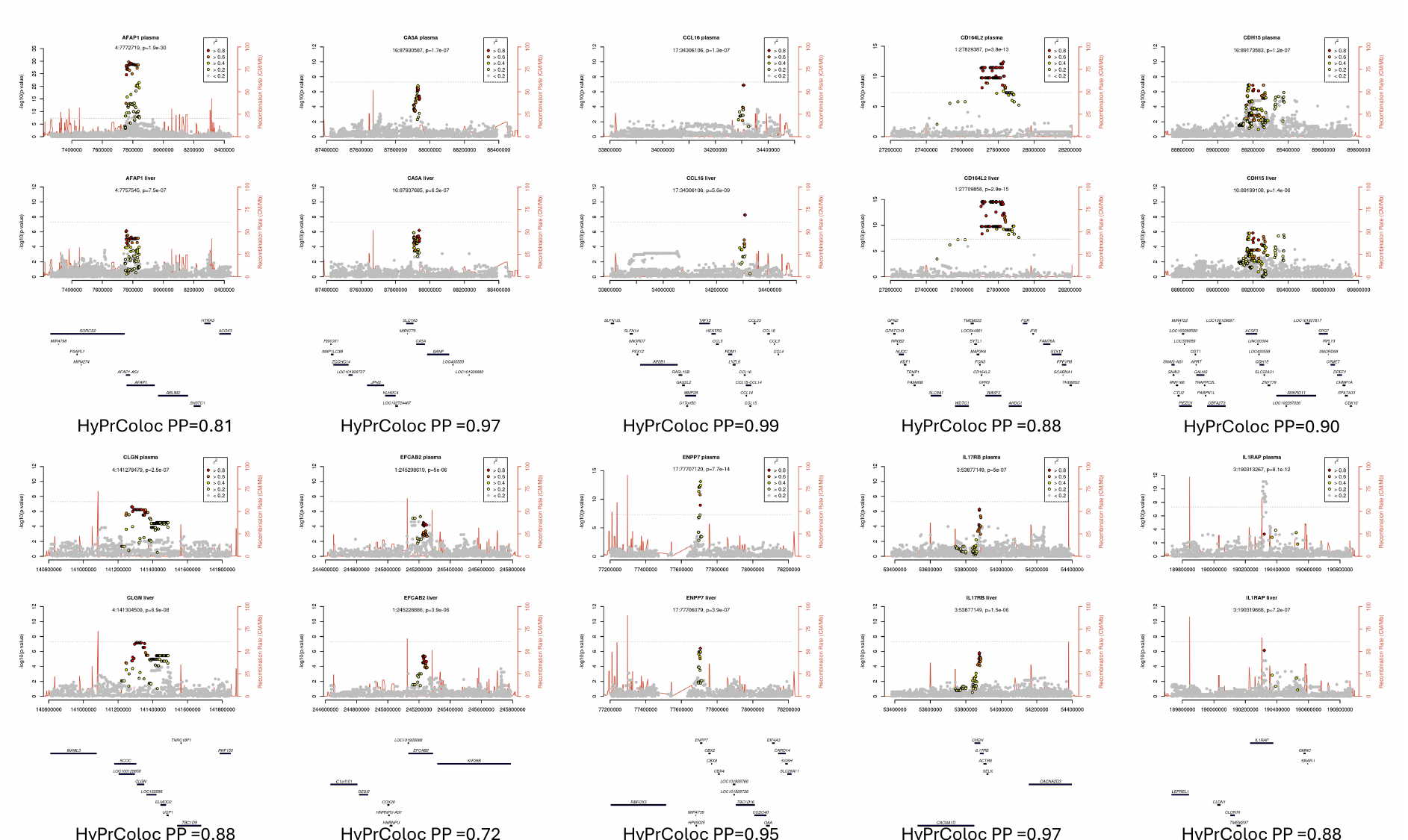

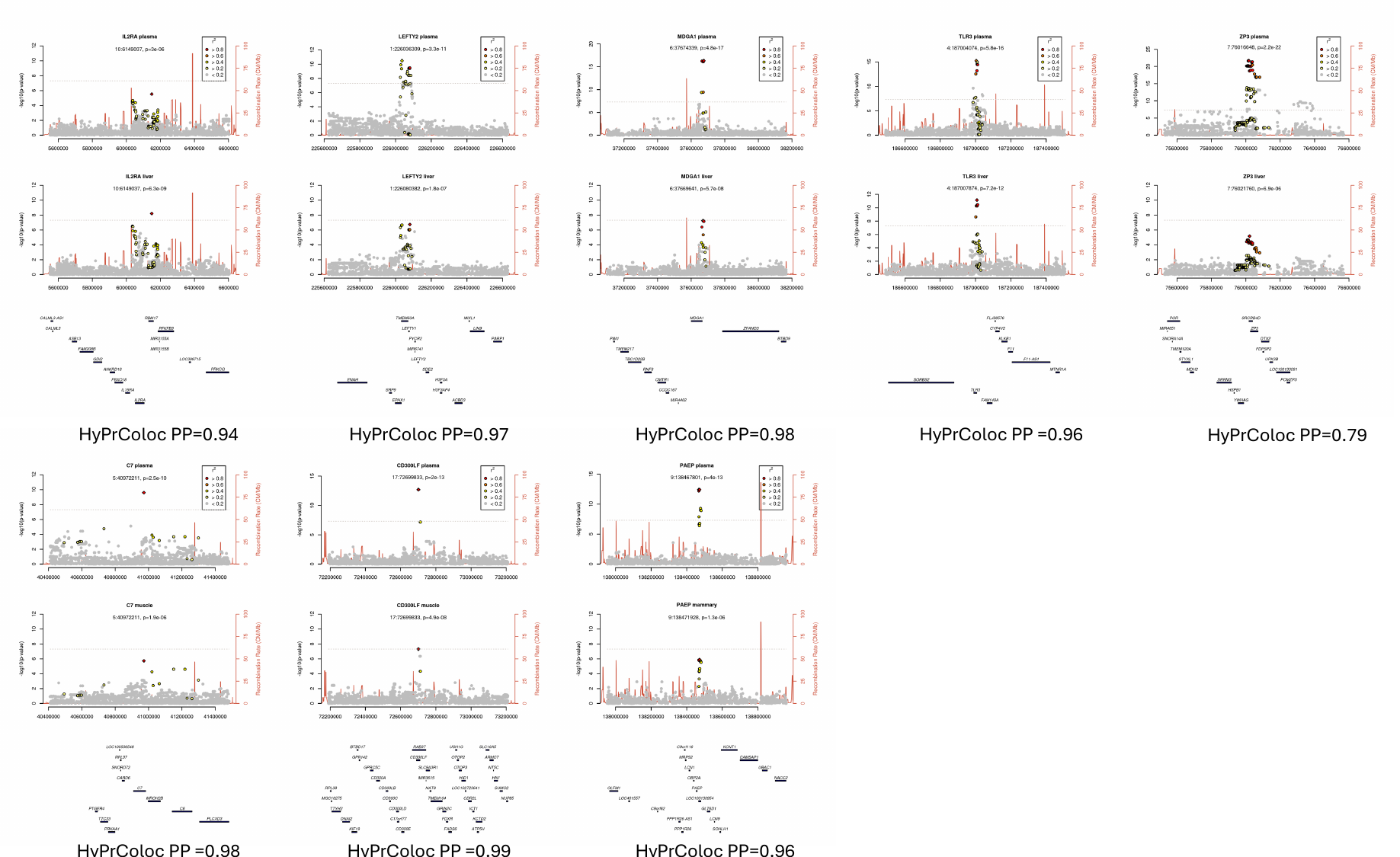

*Supplementary Figure 10: Examples where tissue-specific pQTLs map to tissue specific regulatory regions*

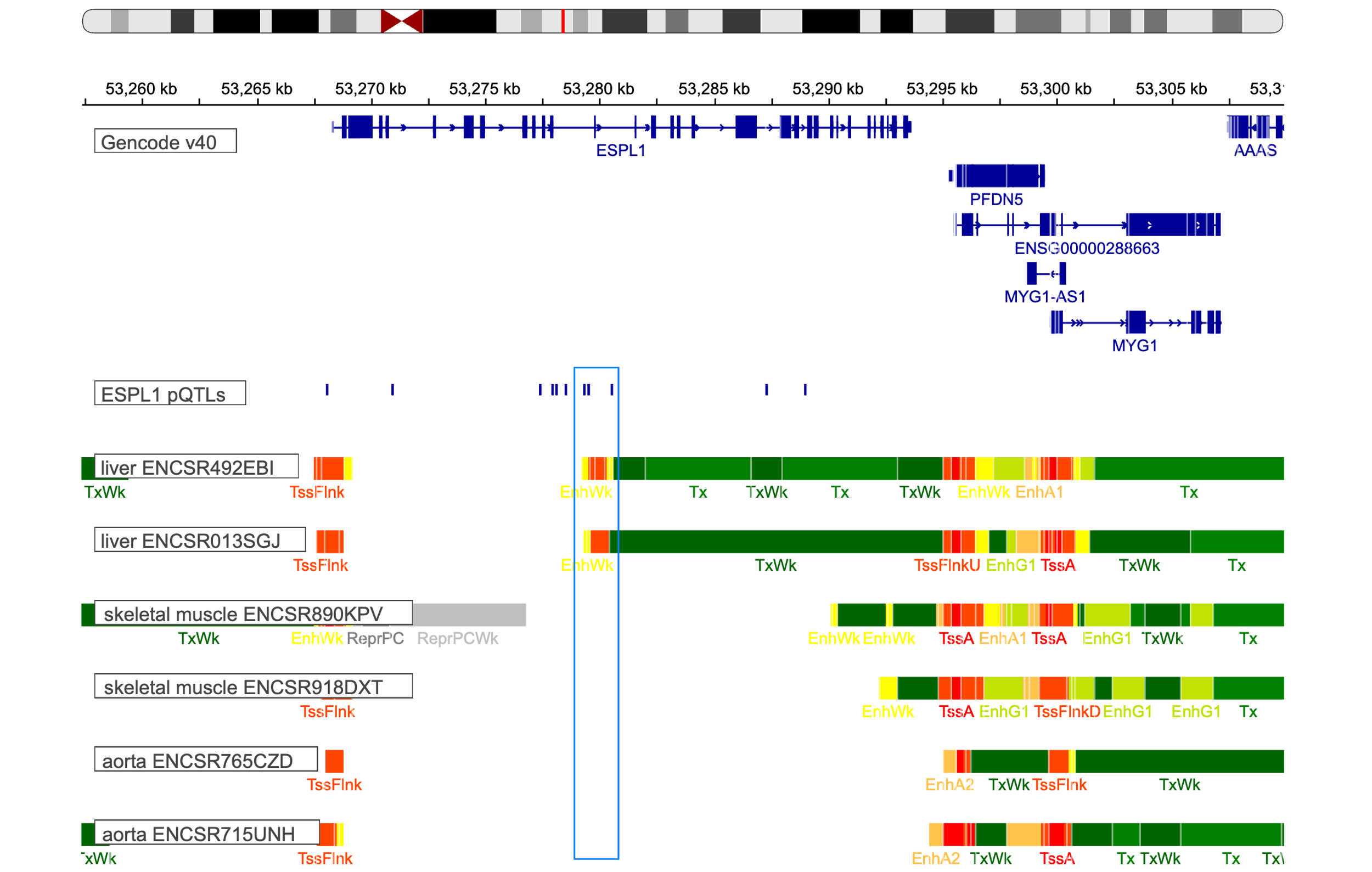

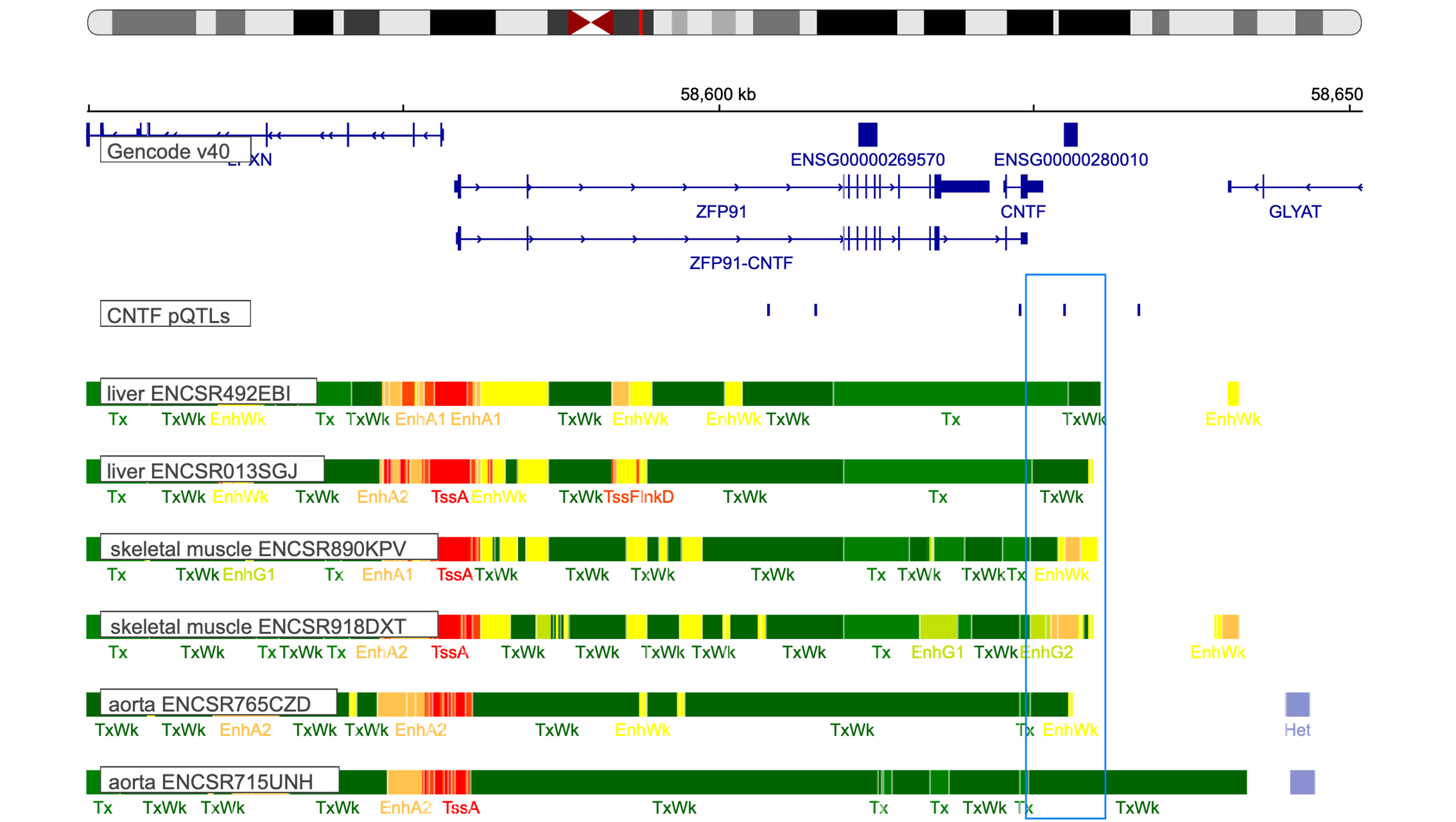

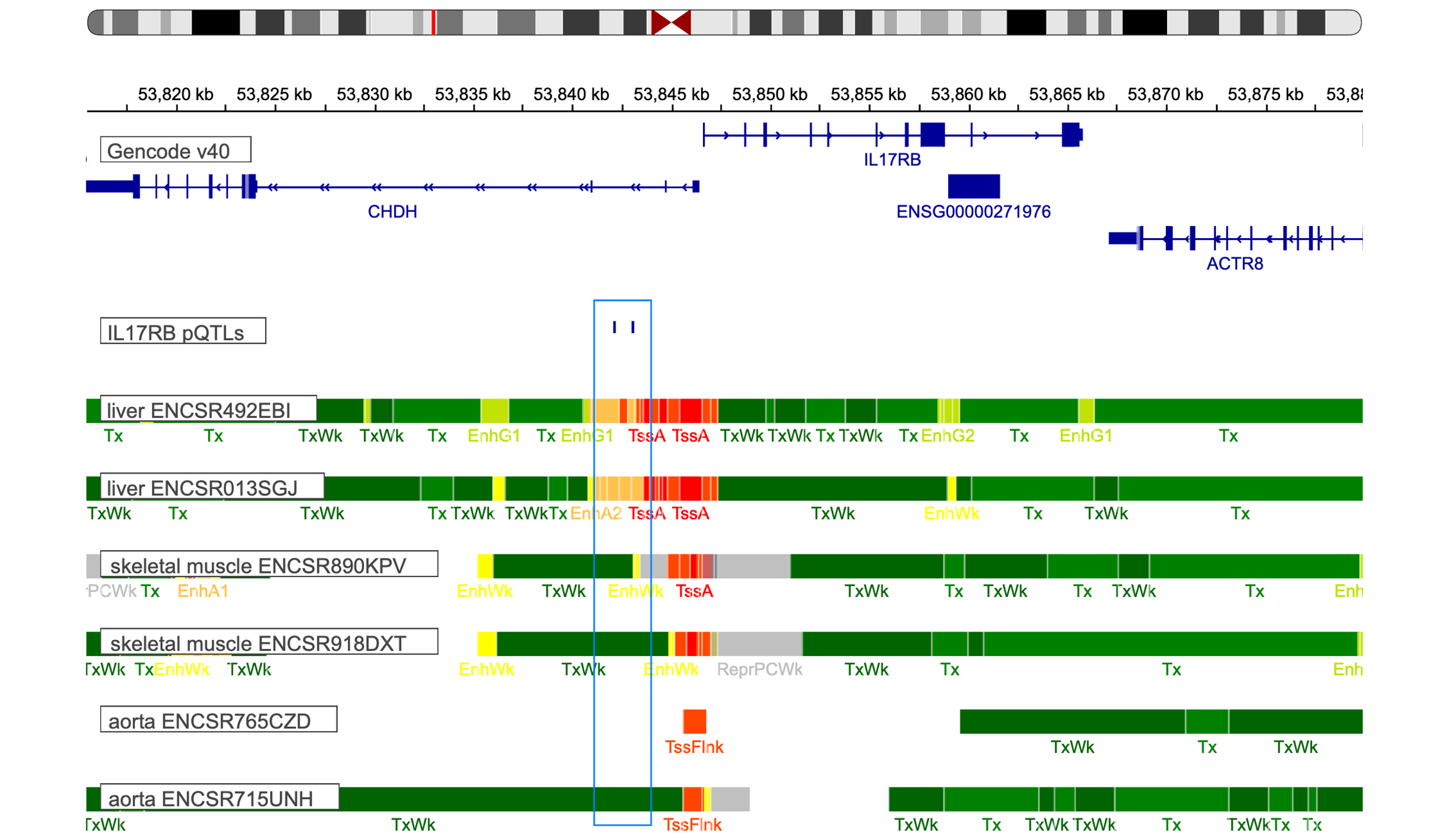

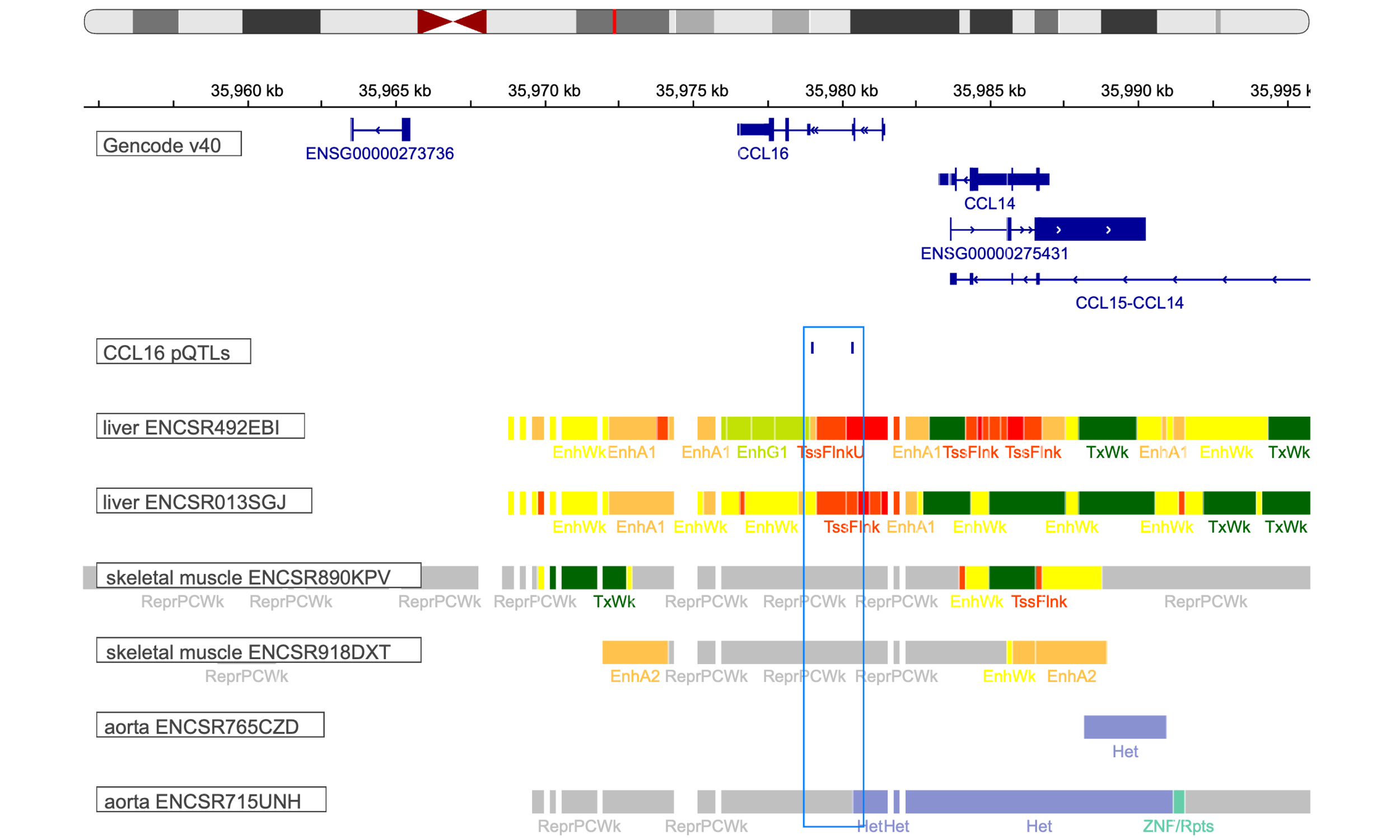

*Chromatin states from ENCODE in liver (dataset identifiers: ENCSR013SGJ and ENCSR492EBI), aorta (dataset identifiers: ENCSR715UNH and ENCSR765CZD) and skeletal muscle (dataset identifiers: ENCSR918DXT and ENCSR890KPV) are plotted with IGV web app (1,2). Regulatory elements are coloured in red, orange and lime green.*

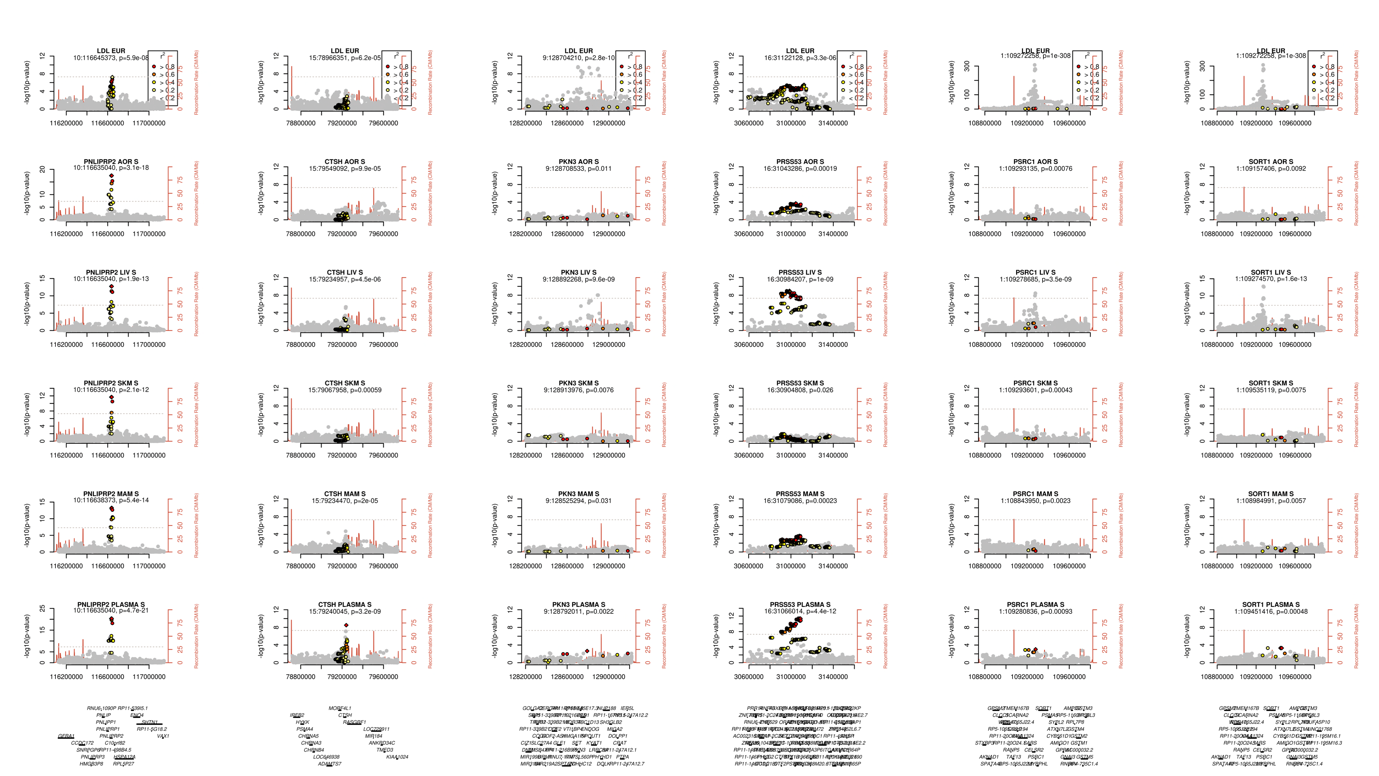
*Supplementary Figure 11: Regional association plots of proteins with tissue pQTLs colocalizing with cardiometabolic traits*

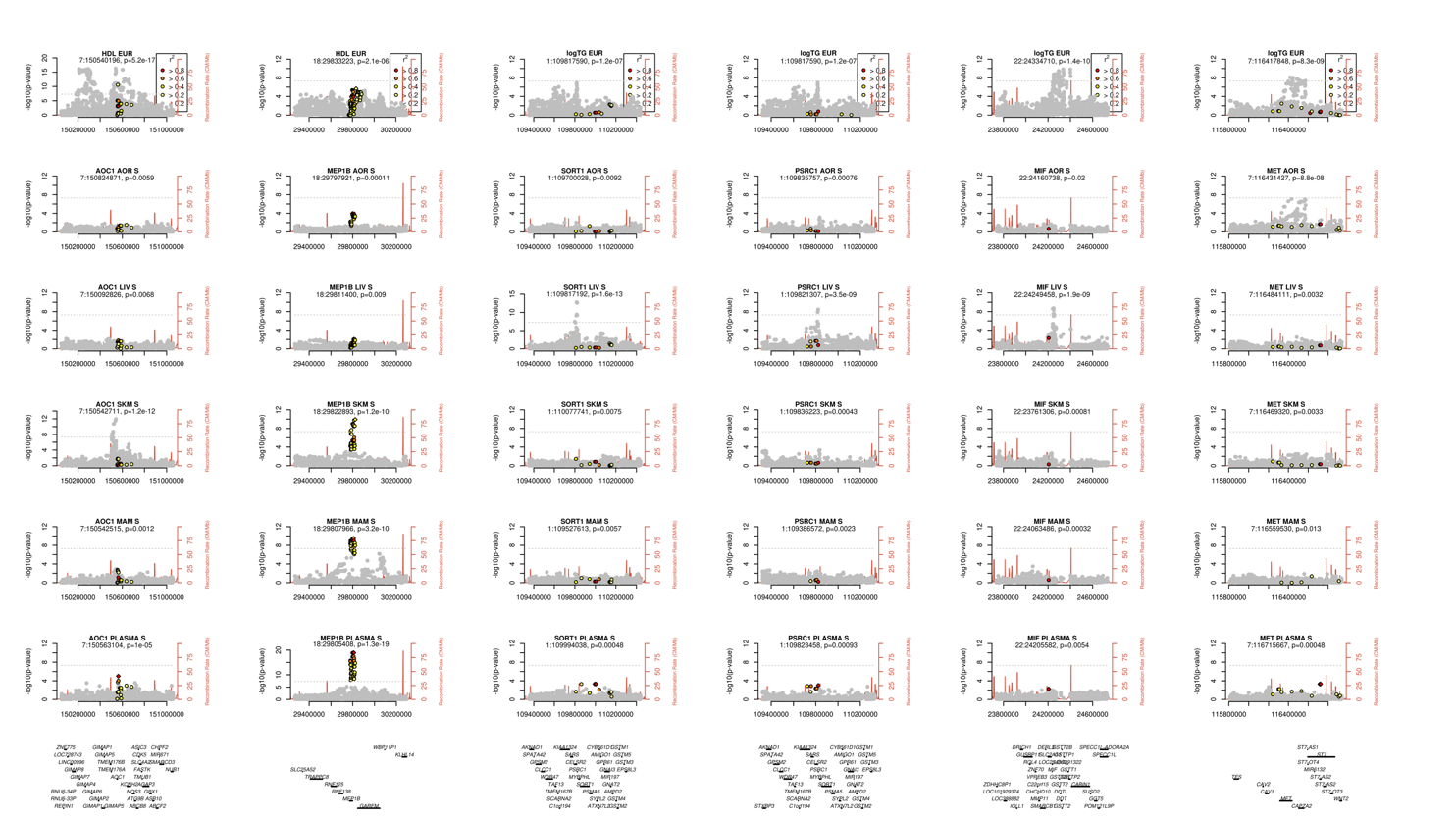

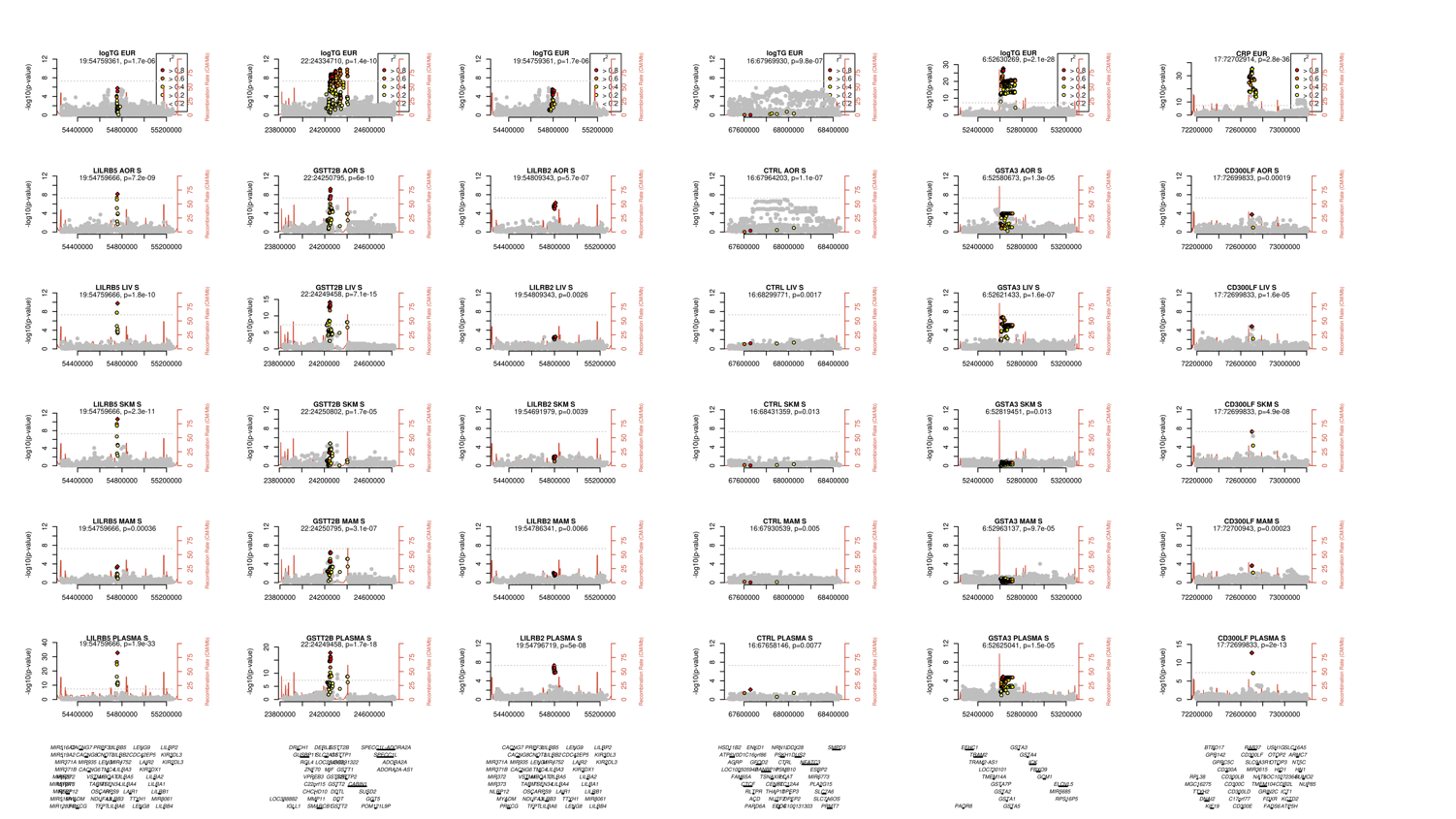

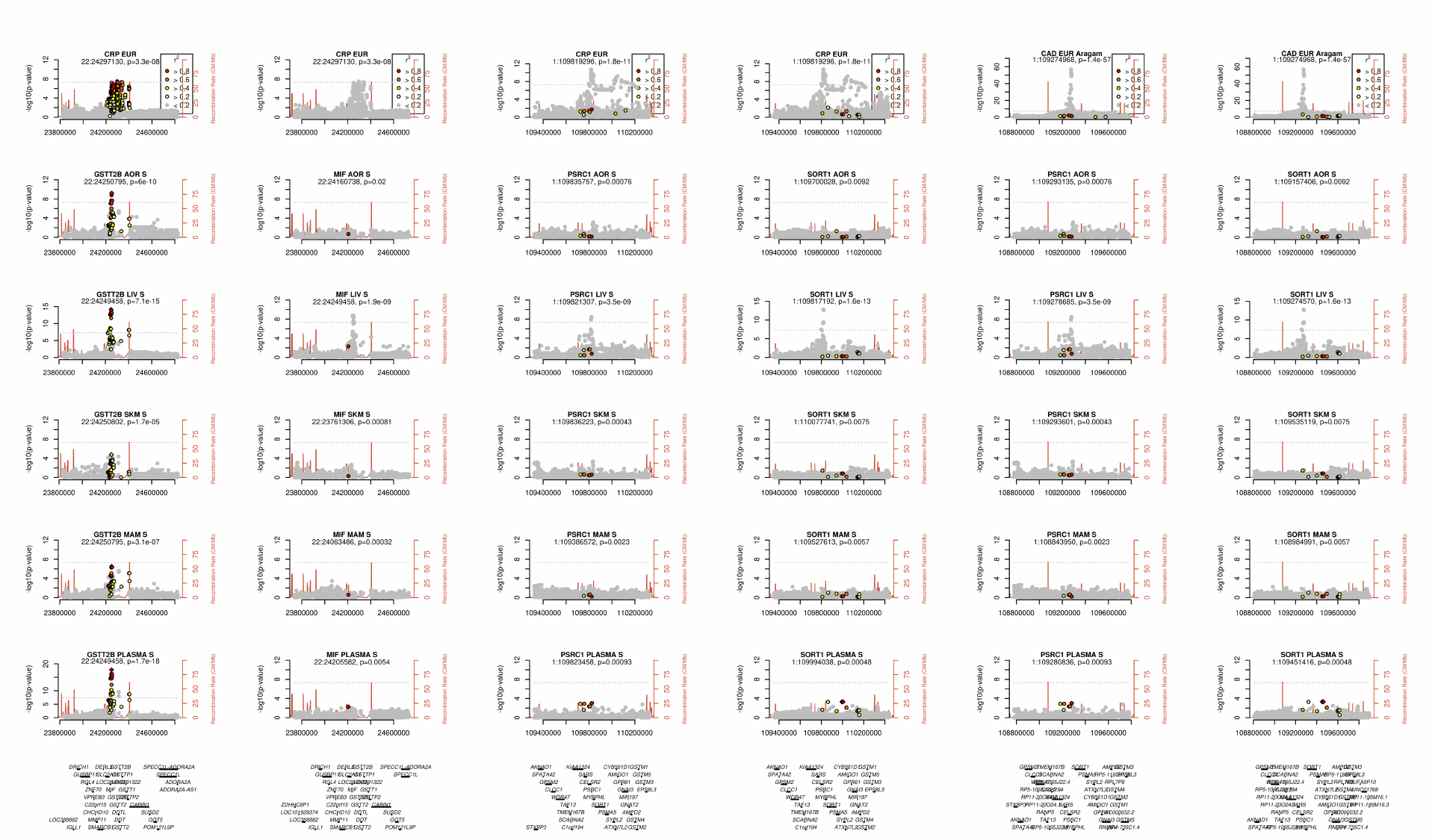

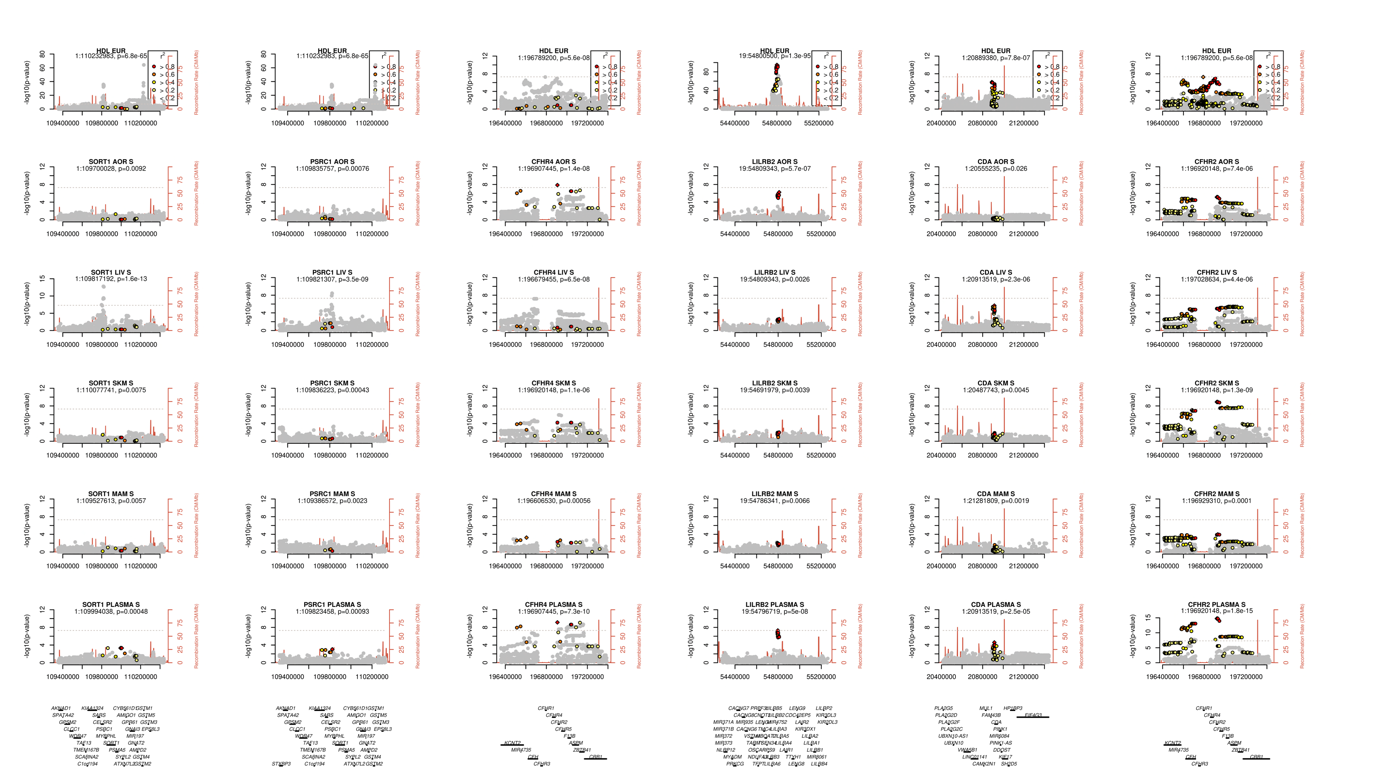

*Supplementary Figure 12: Mendelian randomization results for the effect of increasing protein levels on plasma LDL, HDL, triglyceride and CRP levels*

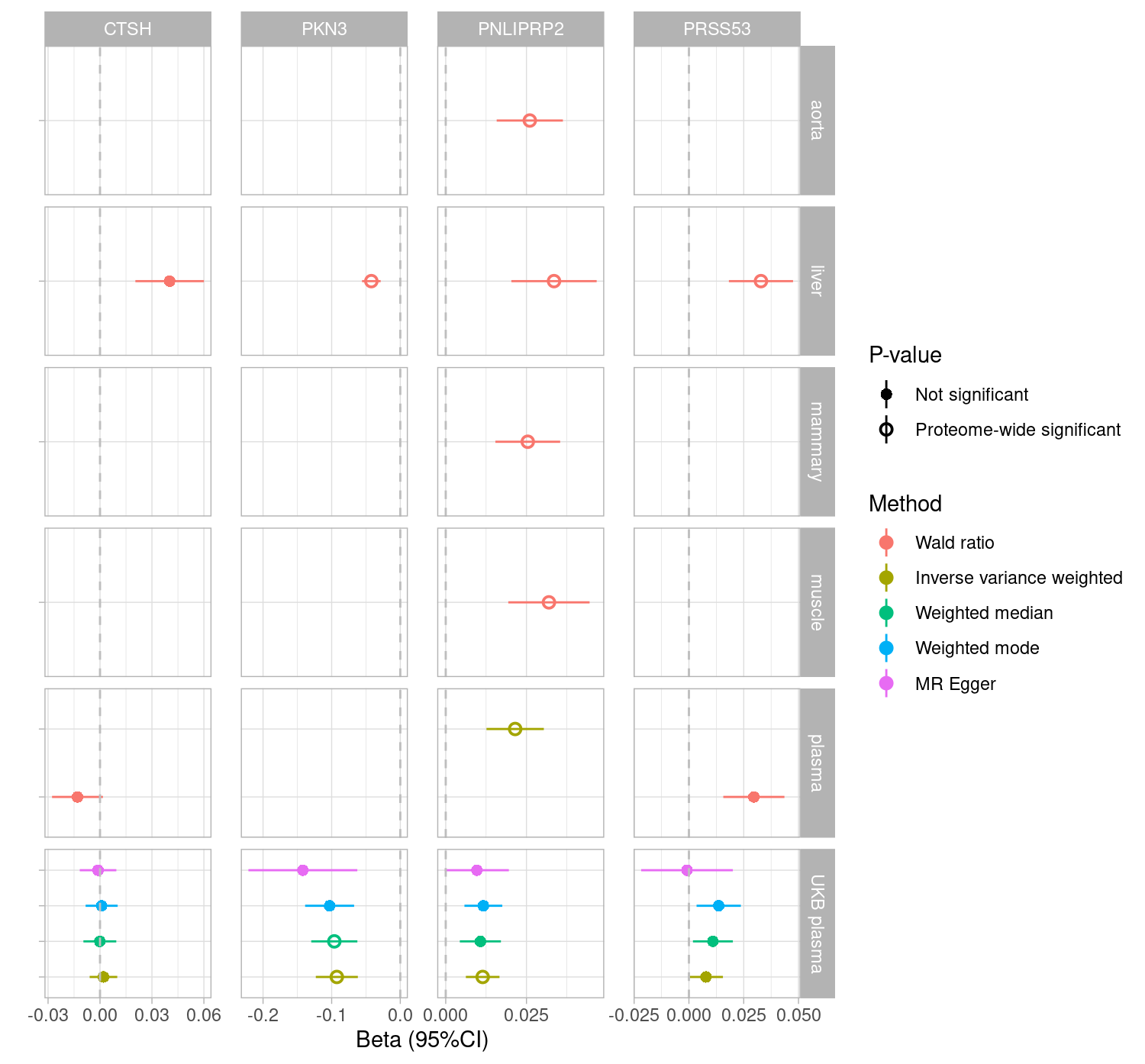

**Outcome=LDL**

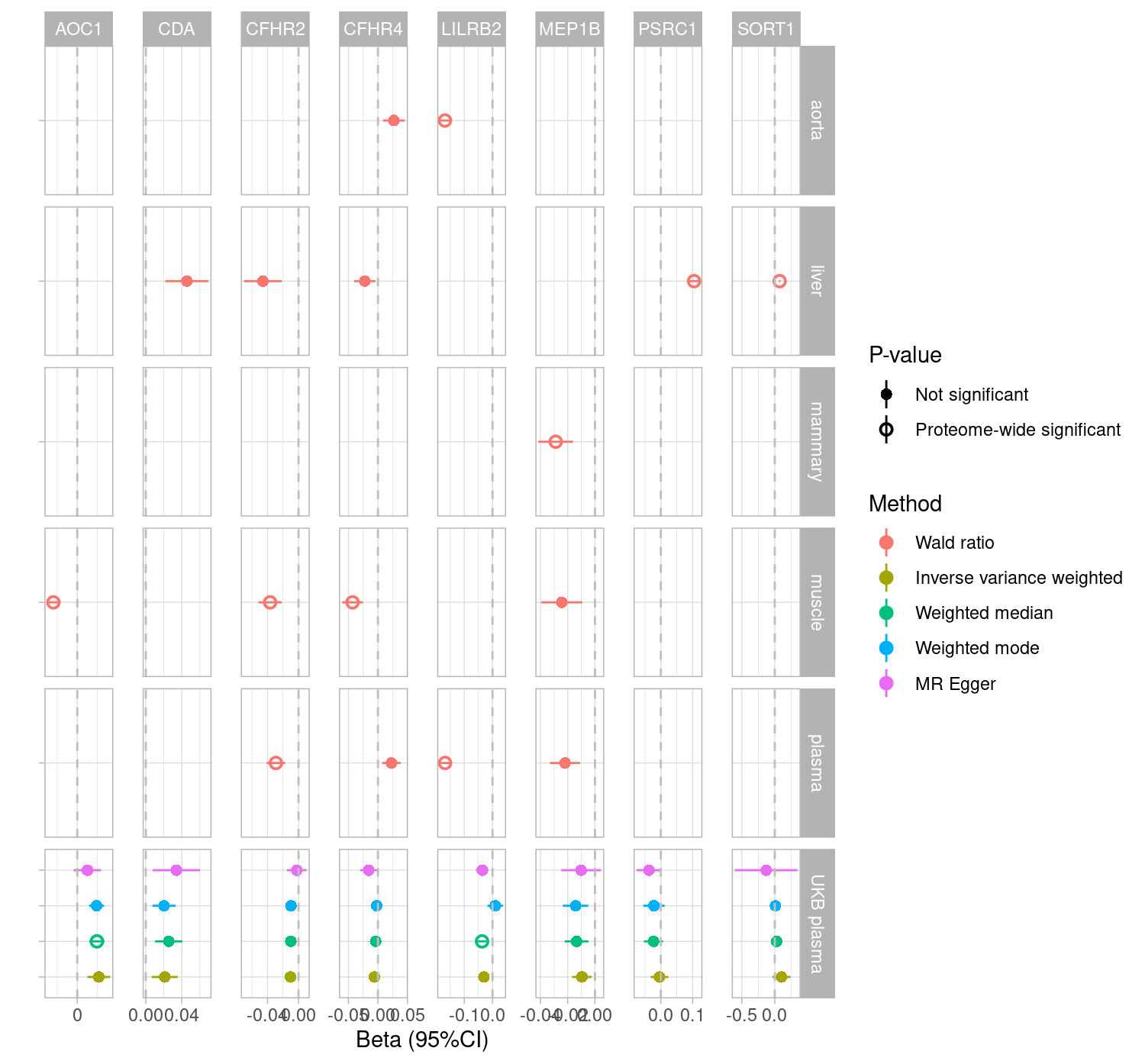

**Outcome=HDL**

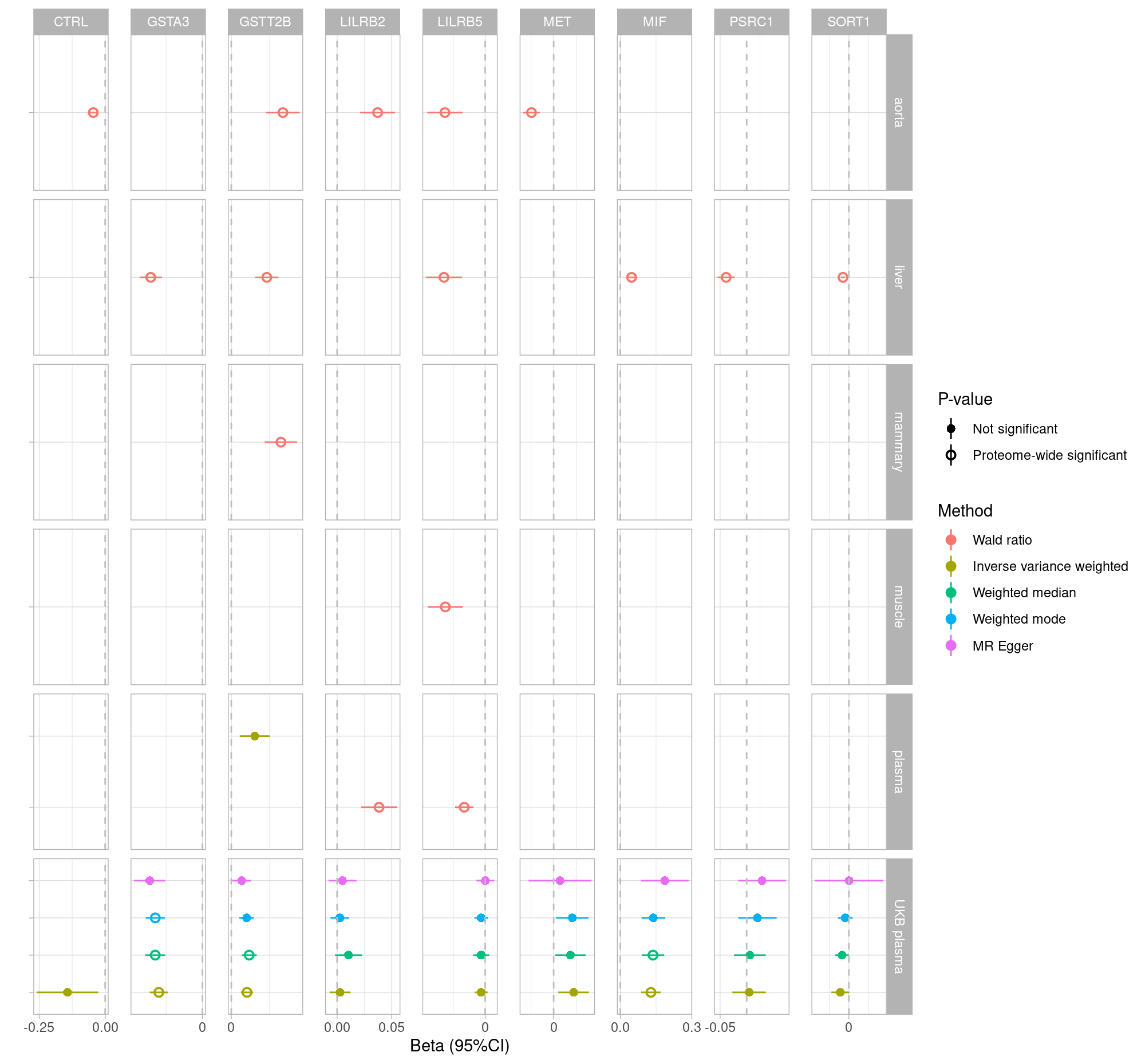

**Outcome=**

**triglycerides**

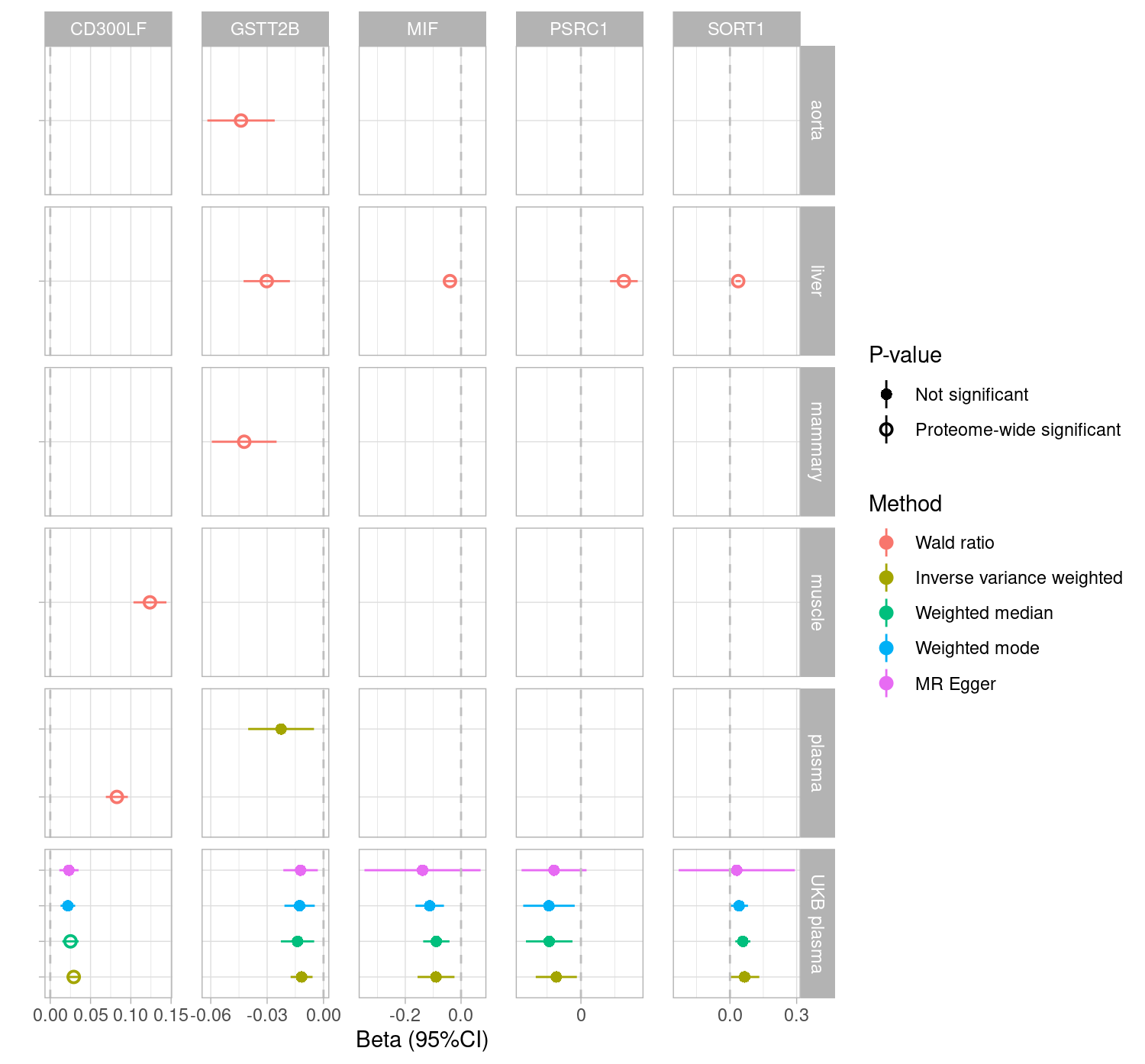

**Outcome=CRP**

*Abbreviations: LDL: low density lipoprotein, HDL: high density lipoprotein, CRP: c-reactive protein.*

*Proteome wide significant: p-value≤0.05/2859 proteins*

*Supplementary Figure 13: Volcano plots of PheWAS results for pQTLs found in tissue but not blood*

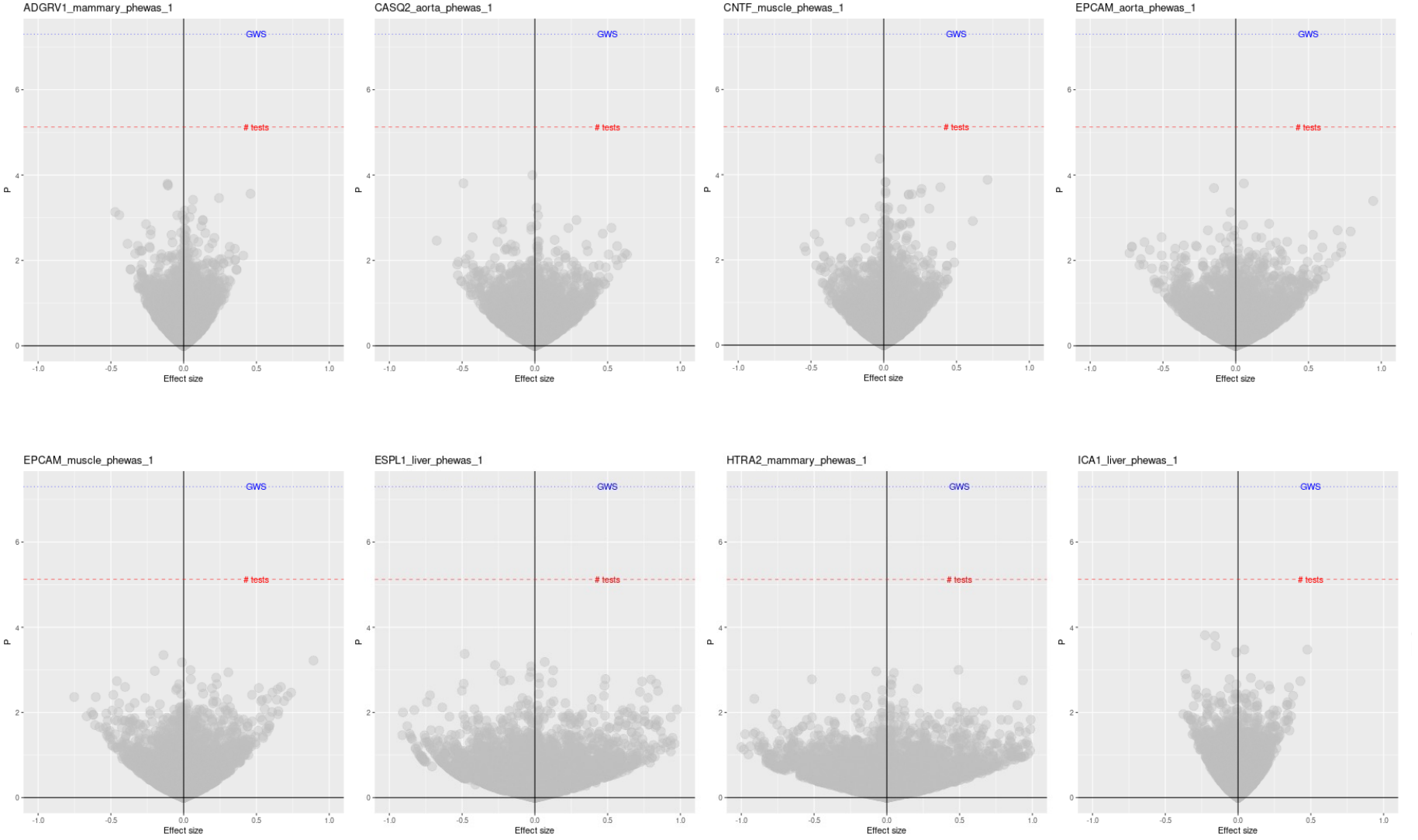

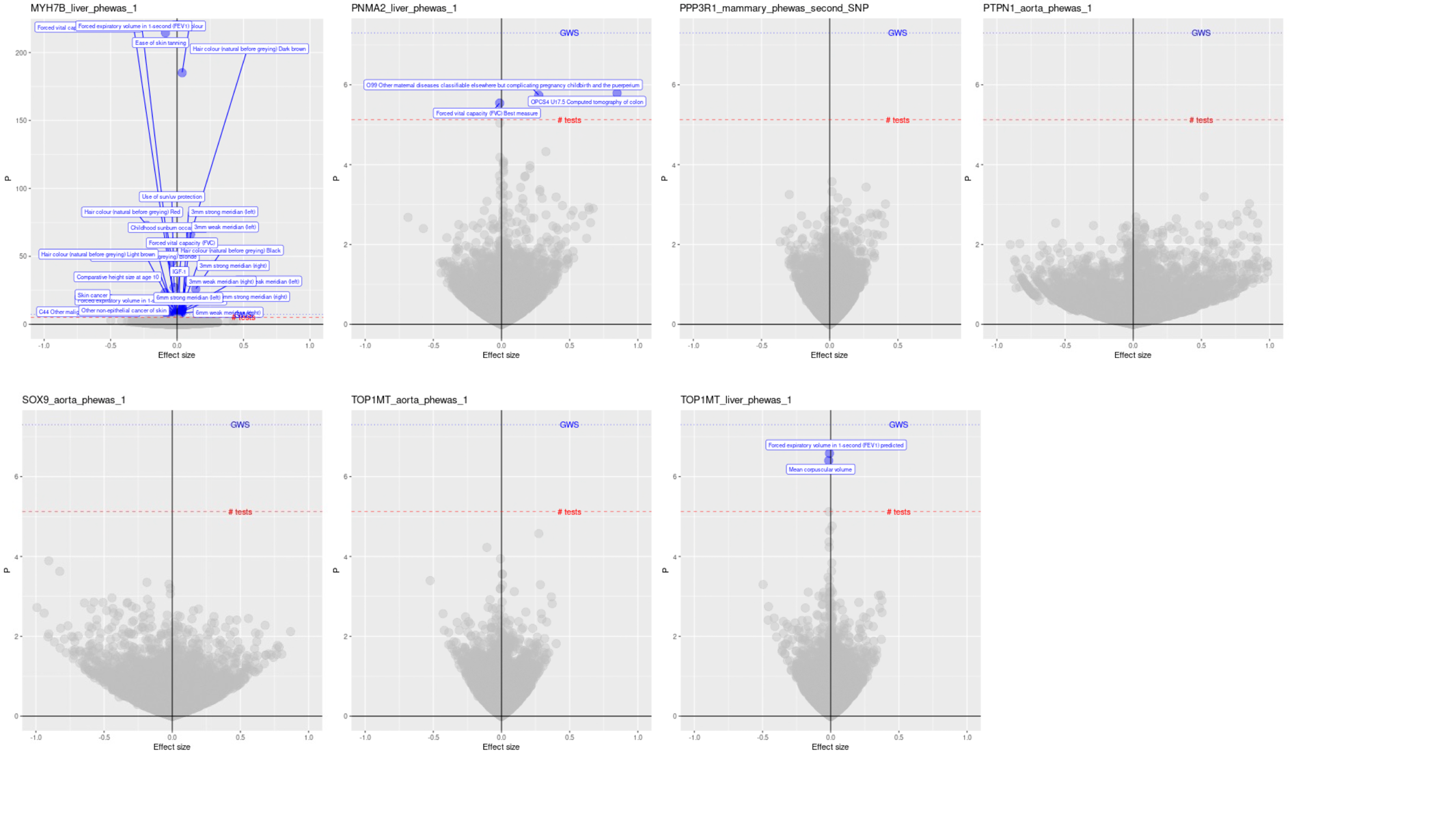

*Supplementary Figure 14: Locus plot of the MYH7B locus and the traits identified as associated with the top liver SNP by PheWAS*

*Supplementary Figure 15: (a) P-values for trans associations of sentinel SNPs at tissue-specific cis-pQTLs with other proteins in blood, (b) local Manhattan plots for these cis-trans protein pairs and (c) Mendelian randomization results for the causal effects of tissue protein on blood protein*
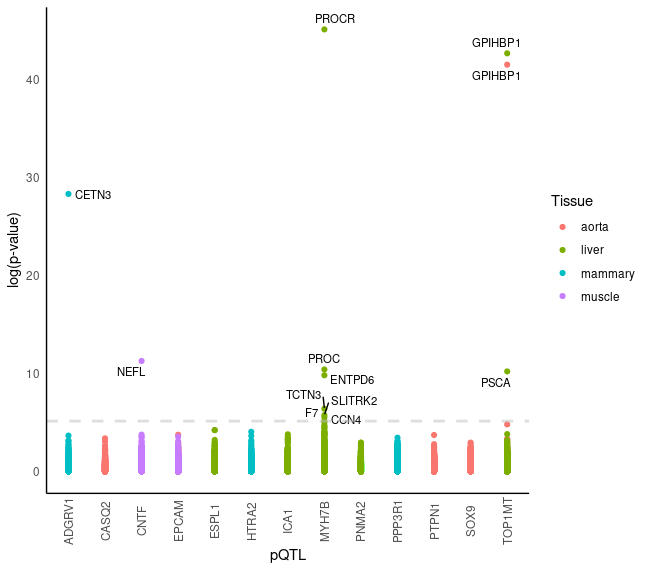

(a)

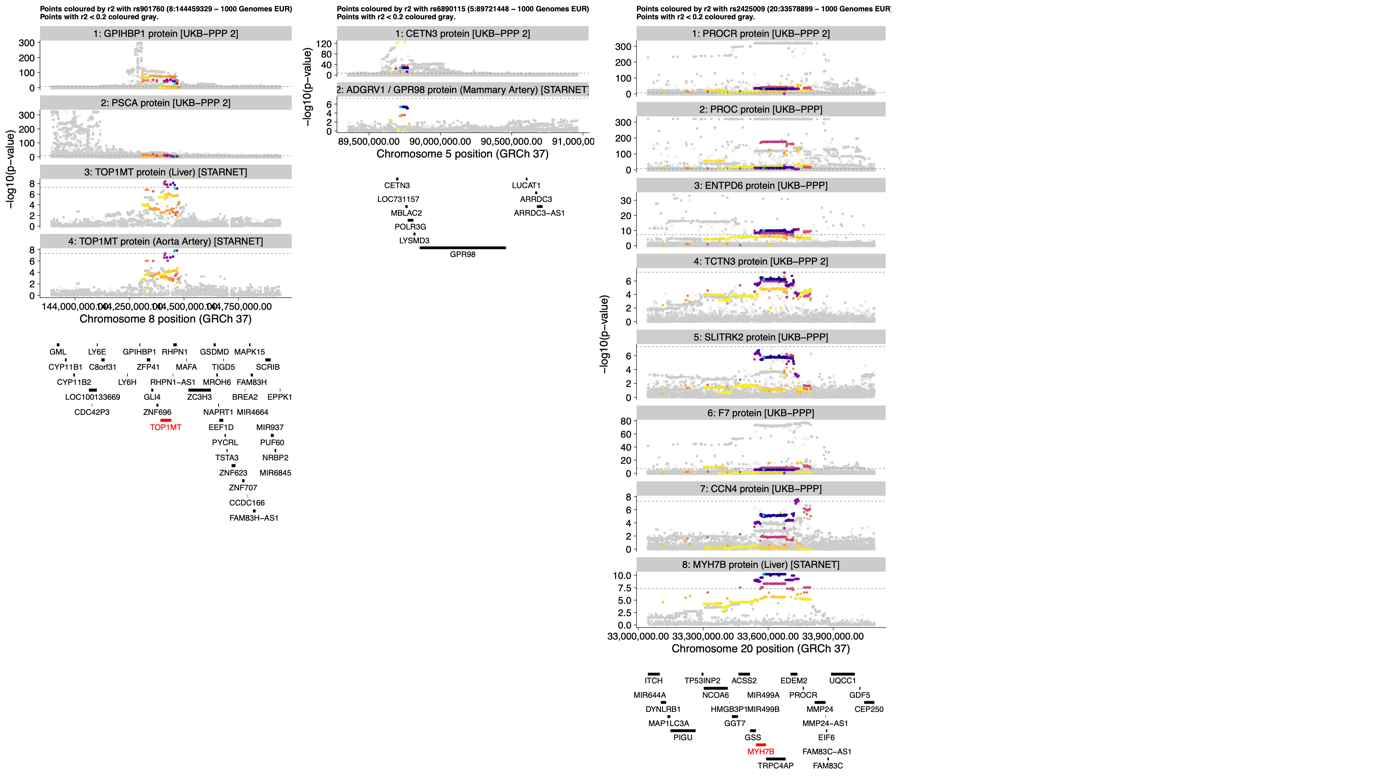

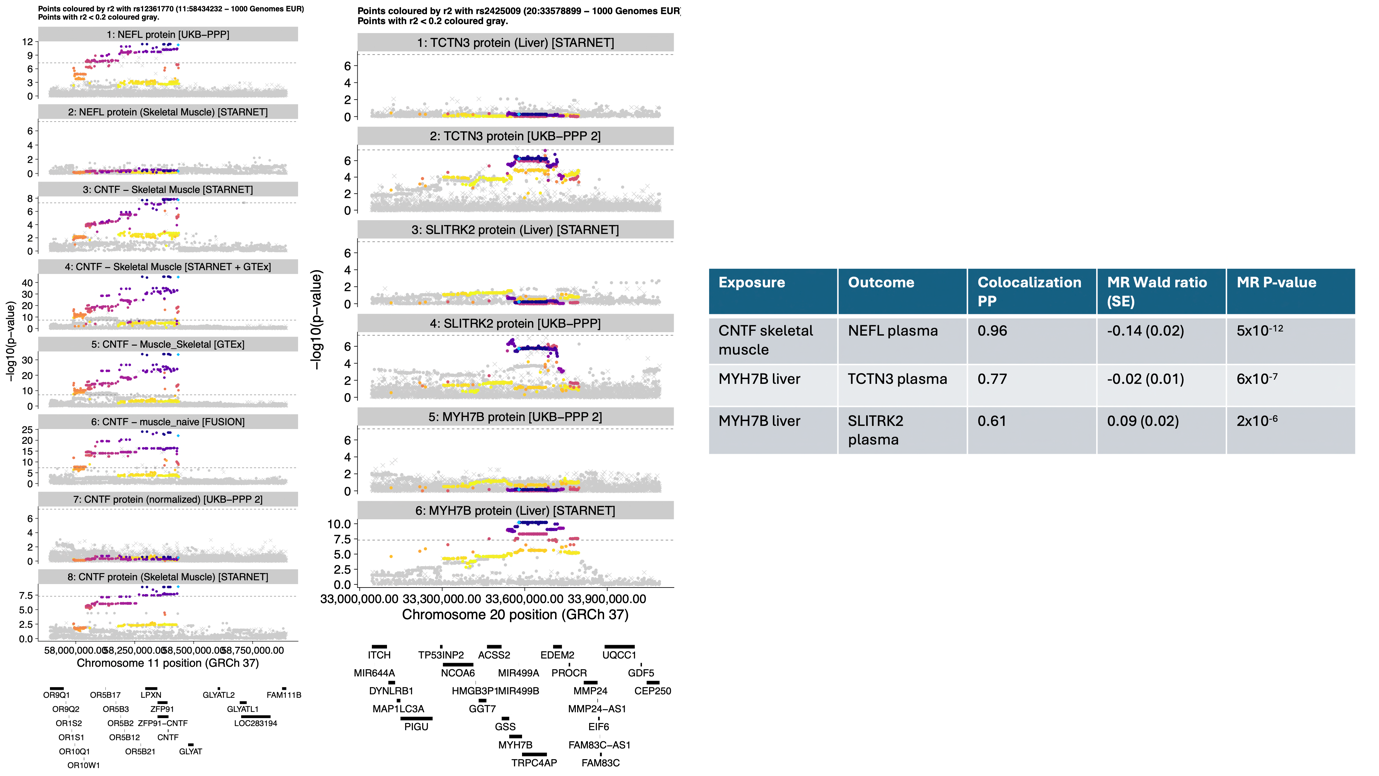

(b)

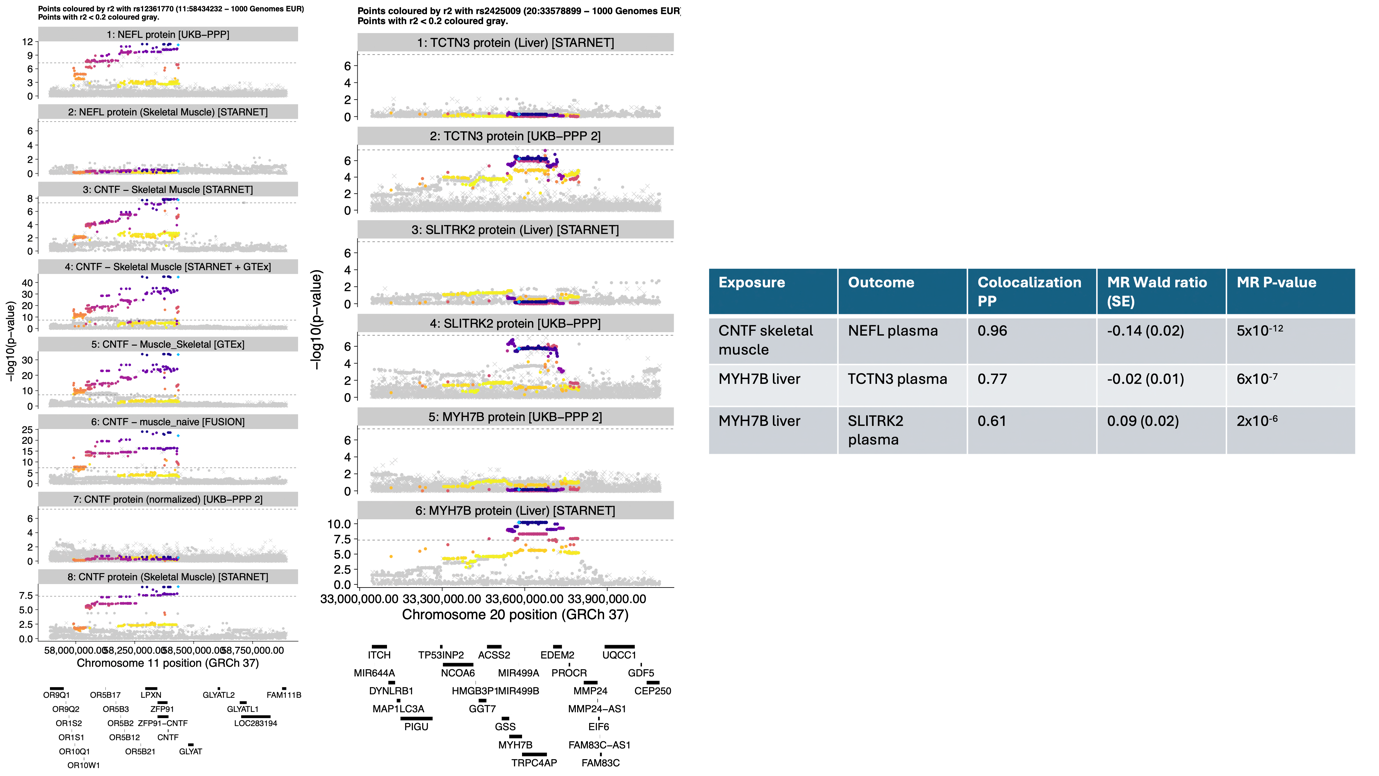

(c)

*Supplementary Figure 16: Power calculation for trans-pQTLs based on an alpha value of 5x10^-8^*

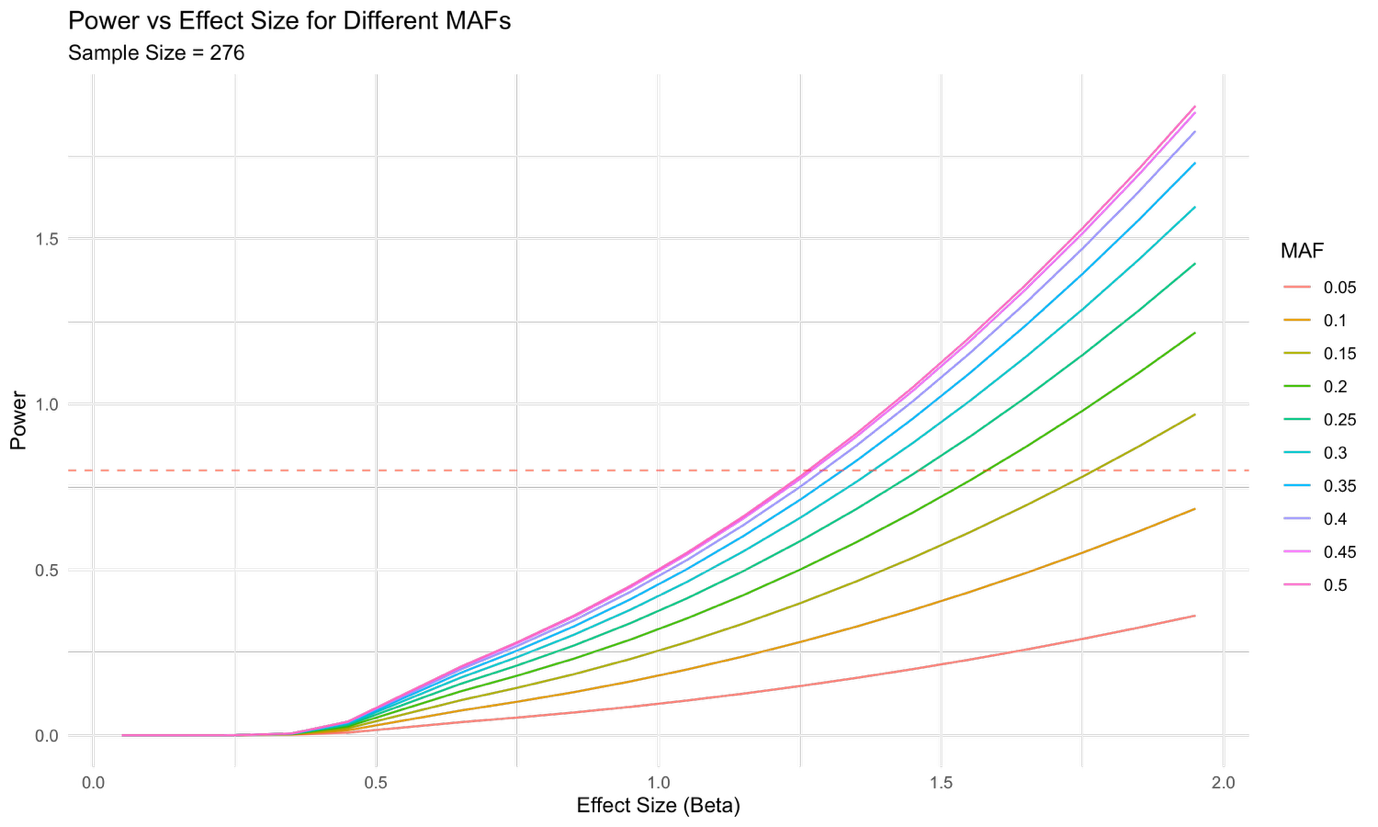

*Abbreviations: MAF: Minor Allele Frequency. Power calculations performed using the R “pwr” package*
